# Spatial gene expression at single-cell resolution from histology using deep learning with GHIST

**DOI:** 10.1101/2024.07.02.601790

**Authors:** Xiaohang Fu, Yue Cao, Beilei Bian, Chuhan Wang, Dinny Graham, Nirmala Pathmanathan, Ellis Patrick, Jinman Kim, Jean YH Yang

## Abstract

The increased use of spatially resolved transcriptomics provides new biological insights into disease mechanisms. However, the high cost and complexity of these methods are barriers to broad clinical adoption. Consequently, methods have been created to predict spot-based gene expression from routinely-collected histology images. Recent benchmarking showed that current methodologies have limited accuracy and spatial resolution, constraining translational capacity. Here, we introduce GHIST, a deep learning-based framework that predicts spatial gene expression at single-cell resolution by leveraging subcellular spatial transcriptomics and synergistic relationships between multiple layers of biological information. We validated GHIST using public datasets and The Cancer Genome Atlas data, demonstrating its flexibility across different spatial resolutions and superior performance. Our results underscore the utility of *in silico* generation of single-cell spatial gene expression measurements and the capacity to enrich existing datasets with a spatially resolved omics modality, paving the way for scalable multi-omics analysis and new biomarker discoveries.

## Introduction

Spatially resolved transcriptomics (SRT) profiling technologies provide spatially resolved mapping of gene expression, and hold the potential to revolutionise our understanding of multicellular biological systems. SRT can enable new insights to improve the prediction of key clinical outcomes of complex diseases, including cancers^1–4^. The recent subcellular *in situ* imaging-based SRT, herein referred to as subcellular spatial transcriptomics (SST), platforms (e.g., 10x Xenium^5^; NanoString CosMx^6^; Vizgen MERSCOPE) can map spatial gene expression at subcellular resolution. Spot-based SRT platforms (e.g. 10x Visium) typically capture the gene expressions of multiple cells per spot, generating spot-based spatial gene expression (SGE). In contrast, the newer SST data are assayed at a resolution smaller than the typical cell and are typically passed through a cell segmentation workflow to provide “single-cell SGE”. This finer spatial resolution of gene expression unlocks the capacity to explore a variety of hypotheses that were previously inaccessible and provide deeper biological insights.

Despite the breakthrough capability in SRT, the high cost associated with these technologies hinders their broad clinical adoption. On the other hand, histopathology images, such as haematoxylin-and-eosin (H&E) stained images, are routinely collected and widely available. H&E-stained images contain rich visual details on cellular and tissue structure that reflect the integration of the underlying biological information in the tissue^7^. The capacity to predict SGE from histopathology alone would unlock insights into variation in gene expression and cell type specific expression that were previously unattainable from the large library of H&E images available. This will open up a new pathway to the discovery of novel biomarkers and therapeutic targets for complex diseases.

Deep learning (DL)-based methods have been developed to facilitate the prediction of spot-level spatial profiles of gene expression values from H&E-stained images^8–14^. They include ST-Net, Hist2ST, HisToGene, DeepPT, and BLEEP. These methods learn visual and spatial associations between matched H&E images and spot-level SGE data. Existing methods typically examine H&E image patches with a Convolutional Neural Network (CNN)^8,10,12^ or Transformer^9,11^ backbone. For example, ST-Net leveraged a CNN-based backbone to extract an embedding of image patches that corresponded to spots, then applied a fully connected layer to predict gene expressions of each spot. Some methods further examined the relationships between adjacent spots on a more global level, for example with Graph Neural Networks (GNNs). For example, Hist2ST combined a Convmixer, Transformer, and a GNN to extract patch-based features, and considered the spatial relationships among the spots to enhance spatial dependencies from the entire image. Furthermore, an alternative strategy is to apply super-resolution to spot-based data^14^. Super-resolved pixels, however, potentially contain a mixture of cells with background areas, and are thus not at true single-cell resolution. Our previous benchmarking study^15^ on the existing methods revealed a weak translational potential, which is likely due to the limited information in the predictions. In these results, expressions in each spot describe a mixture of cells and a mixture of cell types, which contain limited power as the spot-based SGE expressions are not at single-cell resolution. None of these methods have formulated the spatial associations to leverage subcellular resolution SGE data from newer platforms such as 10x Xenium. Methods have been developed to leverage histology and SGE data to predict high-resolution SGE, however, they require input spot-based SGE data during prediction, thereby limiting wider applicability.^16–18^

The increase in resolution from spot-level to single-cell level poses a substantial challenge, as this presents an increased number of expression profiles (from hundreds to thousands of spots, to over tens of thousands of cells), and also necessitates the ability to discern finer-grained variations in H&E images that correspond to subtle variations in gene expression. These fine-grained details in H&E images are typically noisy and may provide limited information for discerning patterns that are even broader than gene expression, such as cell types. Different cell types that express distinct gene markers and typically exhibit similar morphological features in histology may be difficult to tell apart with morphological information alone, e.g., some types of lymphocytes including B and T cells. Additionally, gene expression within a target cell is known to be influenced by neighbouring cell types, for example, various fibroblast subtypes that occur in the presence of macrophages or malignant cells. Overall, there are complex relationships between the histological appearance of cells, gene expression within cells, cell type, and composition of neighbouring cell types. A DL model can be designed to leverage these relationships to achieve better gene expression predictions. To our knowledge, there is not yet a method that is specifically designed for single-cell SGE prediction from histology images, and the relationships between these multiple layers of biological information into such a method remains to be leveraged.

Here, we introduce a novel framework to predict spatially resolved single-cell **G**ene expression from **HIST**ology, **GHIST** (pronounced d3Ist; **Figure 1**). GHIST addresses the challenges of predicting single-cell SGE from histology images through a number of key innovations. Most importantly, GHIST leverages multiple prediction components that use the relationships between multiple layers of information (including cell type, neighbourhood composition, nuclei morphology, single-cell expressions). GHIST synergistically captures interdependencies between information types by its multitask architecture and multiple loss functions, which overall improves SGE prediction. Furthermore, GHIST is designed to handle different resolutions in addition to single-cell-level SGE, such as spot-based SGE, for which we demonstrate the superior performance of GHIST compared to the state-of-the-art approaches. Along with the development of our method, we demonstrate the power of GHIST in providing an innovative *in silico* strategy to add a layer of rich spatial information to existing large-scale databases, without costly practical experiments. This can facilitate the generation of new hypotheses for spatial experiments, the discovery of cell-type-specific and spatial associations between biomarkers, and relationships that provide new insights into disease mechanisms.

**Figure 1.**
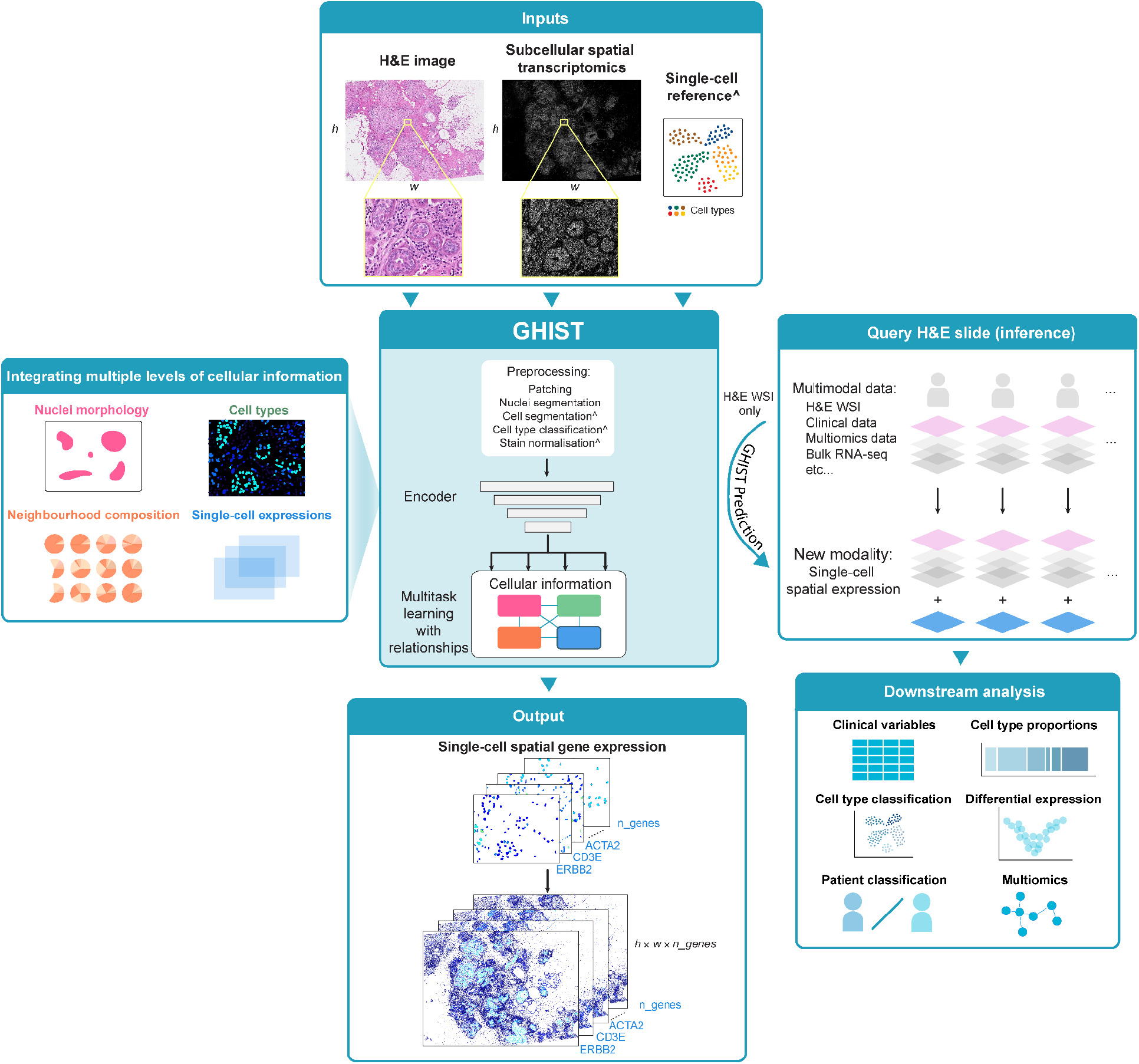
GHIST framework. GHIST maps H&E images to spatially resolved single-cell gene expressions through leveraging paired SST data during training. The multitask DL-based model integrates relationships between multiple levels of information related to gene expressions within cells. Once trained, GHIST enables the *in silico* estimation of spatially resolved single-cell gene expressions from existing H&E data alone, that can be used to enrich further downstream analysis. The cell segmentation and cell type classification preprocessing steps are performed during training only. During inference, stain normalisation is applied on query H&E slides, and single-cell reference information is not required as an input if it was not used during training.

## Results

### GHIST: Leveraging multi-level biological information enables spatially resolved single-cell gene expression (SGE) prediction

GHIST is a multitask DL-based learning method that enables spatially resolved SGE prediction at single-cell-level resolution by mapping H&E images to the expression of hundreds of genes within individual cells in the image, and providing spatial localisation of cells (**Figure 1**). In other words, GHIST maps H&E images to a collection of SGE images (where the number of channels equals the number of genes, and pixel intensities equal the total expression of individual genes within cells). GHIST can achieve this mapping by learning from samples comprising an H&E image and its corresponding SST data. Once trained, SST data are not used during inference of SGE. In other words, the trained model is applied to unseen H&E data only and no additional spatial omics level information is required. During training, single-cell RNA-seq data are leveraged to improve the predicted expression profiles by providing known cell type information, and may be extracted from public repositories such as the Human Cell Atlas or CZI CELLXGENE. There is no requirement for the single-cell data to be matched to the input H&E images.

GHIST considers the interdependencies between four levels of biological information, including (i) cell type, (ii) neighbourhood composition, (iii) cell nucleus morphology, and (iv) single-cell RNA expression. GHIST leverages its multitask DL architecture with four prediction heads (corresponding to the information types) to jointly learn these types of information for each cell. Features and predictions provided by each prediction head are used between different prediction heads, as an input or through training losses that help the model to learn the interdependencies. The design of the prediction heads and loss functions is informed by biological knowledge to capture interdependencies between information types. For example, the composition of the local neighbourhood is used to inform gene expression prediction, and gene expression within cells are used to predict cell type to ensure the expressions are biologically meaningful (details in Methods). This design allows GHIST to elucidate the variations in gene expression within single cells from H&E images, and enables better predictions of expressions for cell types that are difficult to distinguish in histology (see Methods and **Supplementary Figure 1**).

### GHIST effectively captures gene expression at single-cell resolution

We used a range of evaluation metrics including cell type proportion, spatially variable genes (SVGs), and correlation to evaluate the reliability of the gene expressions predicted by GHIST. Here we define the “ground truth” single-cell expressions as those obtained through carrying out cell segmentation on SST data, while the cell types were obtained by applying a standard supervised cell annotation tool, scClassify, on the expressions (details in Methods). **Figure 2a-d** and **Supplementary Figure 2** compare the cell types predicted based on the predicted expressions from GHIST to the ground truth cell types for two breast samples. The cell type distributions on the slides (**Figures 2a and 2c**), including overall cell type composition (**Figures 2b and 2d**), were strikingly similar between the ground truth and the predictions, showing that the predicted gene expression by GHIST successfully maintained cell type information of the samples.

**Figure 2.**
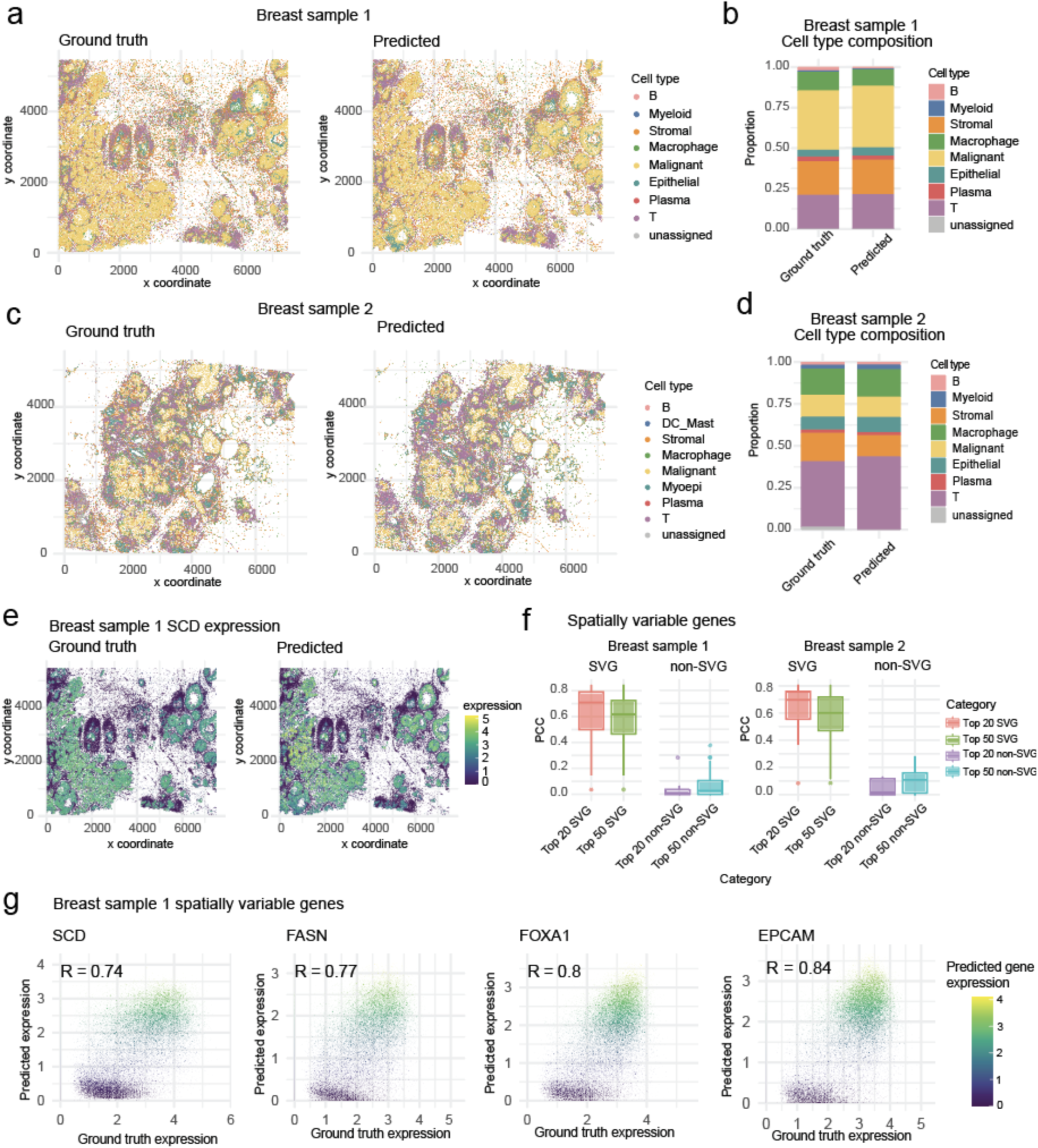
GHIST predictions on two breast cancer H&E images. Comparison between cell types **(a)** and cell type compositions **(b)** obtained from BreastCancer1 paired SST (Xenium) data and predicted cell types from expressions predicted from H&E. Comparison between cell types **(c)** and cell type compositions **(d)** obtained from BreastCancer2 paired SST (Xenium) data and predicted cell types from expressions predicted from H&E. **(e)** Predicted gene expression in individual cells from GHIST and SST data (for the gene *SCD*) on BreastCancer1. **(f)** Boxplot of computed PCC between predicted and measured (ground truth) expressions for top 20 and 50 predicted SVGs and non-SVGs in both BreastCancer1 and BreastCancer2. **(g)** Scatter plots showing the predicted SGE expressions of *SCD, FASN, FOXA1*, and *EPCAM* compared to measured SST expressions.

Next, we examine if the predicted expression values are accurate by examining the correlation between predicted and ground truth expressions amongst spatially variable genes (SVGs) or highly variable genes (HVGs). Using Stearoyl-CoA desaturase 1 (*SCD*) as an example, we observe a strong agreement between the ground truth and predicted gene expression across the slide (**Figure 2e** and **Figure 2f)**. We calculated the correlation between the predicted and measured (ground truth) intensities for the SVGs and the non-SVGs. We found a high correlation between the predicted and measured expression of SVGs, representing biologically meaningful genes, with the top 20 and top 50 SVGs having median correlations of 0.7 and 0.6, respectively (**Figure 2g**). In contrast, as we do not anticipate correlation between two sets of white noise data, genes with low expression should not have correlation between predicted and measured intensities. Consequently, among the non-SVGs, which included many low-expressed genes, we observed a very low correlation of 0 to 0.1, as expected. Some highly correlated SVGs included *SCD* (r = 0.74), *FASN* (r = 0.77), *FOXA1* (r = 0.8) and *EPCAM* (r = 0.84) (**Figure 2h**) which are meaningful genes with known associations with breast cancer^19 20 21 22^. Similar results are observed when we focus on highly variable genes (HVGs) (**Supplementary Figure 3a-b**). Finally, GHIST was applied to two additional datasets - lung adenocarcinoma and melanoma - for which strong agreement between predicted and ground truth cell type proportions is observed (**Supplementary Figures 4 and 5**, melanoma r = 0.92 and lung adenocarcinoma r = 0.97), demonstrating that GHIST is applicable to other cancer tissues.

### GHIST is extendable to spot-based technologies and outperforms state-of-the-art methods

GHIST can be easily adapted to predict spot-based SGE. Here, we use an existing benchmarking framework proposed for spot-base methods^15^ to establish the extensibility of our method. Using the HER2ST dataset^23^, we first evaluated the spot-based SGE predictions with metrics commonly used in the literature such as Pearson Correlation Coefficient (PCC) and Structural Similarity Index (SSIM) across all genes for each method. **Figure 3a** and **3-b** shows that our method achieved the highest PCC of 0.16 (compared to 0.14 for ST-Net and 0.11 for DeepPT), and SSIM of 0.10 (compared to 0.08 for ST-Net and 0.07 for DeepPT). Next, we note that the performance of our method on the top five biologically meaningful genes (**Figure 3c**) had the highest correlation compared to other methods, with *GNAS* (r = 0.42), *FASN* (r = 0.42), *SCD* (r = 0.34), *MYL12B* (r = 0.32), and *CLDN4* (r = 0.32), indicating one should examine correlation among genes with meaningful biological signal. Thus, with the limitation of the PCC values in mind, we examined more meaningful gene characteristics metrics, as discussed in our recent benchmarking work^15^, where we evaluated based on top 10% of HVGs and top 20 SVGs. **Figures 3d and 3e, and Supplementary Figure 6** show that our method provided the highest PCC in HVGs (0.20) and SVGs (0.27), and also SSIM in HVGs (0.17) and SVGs (0.26). Together, these results demonstrate that GHIST predicted SGEs that were more aligned to the ground truth SGE than existing methods and that it improves over existing spot-based methods.

**Figure 3.**
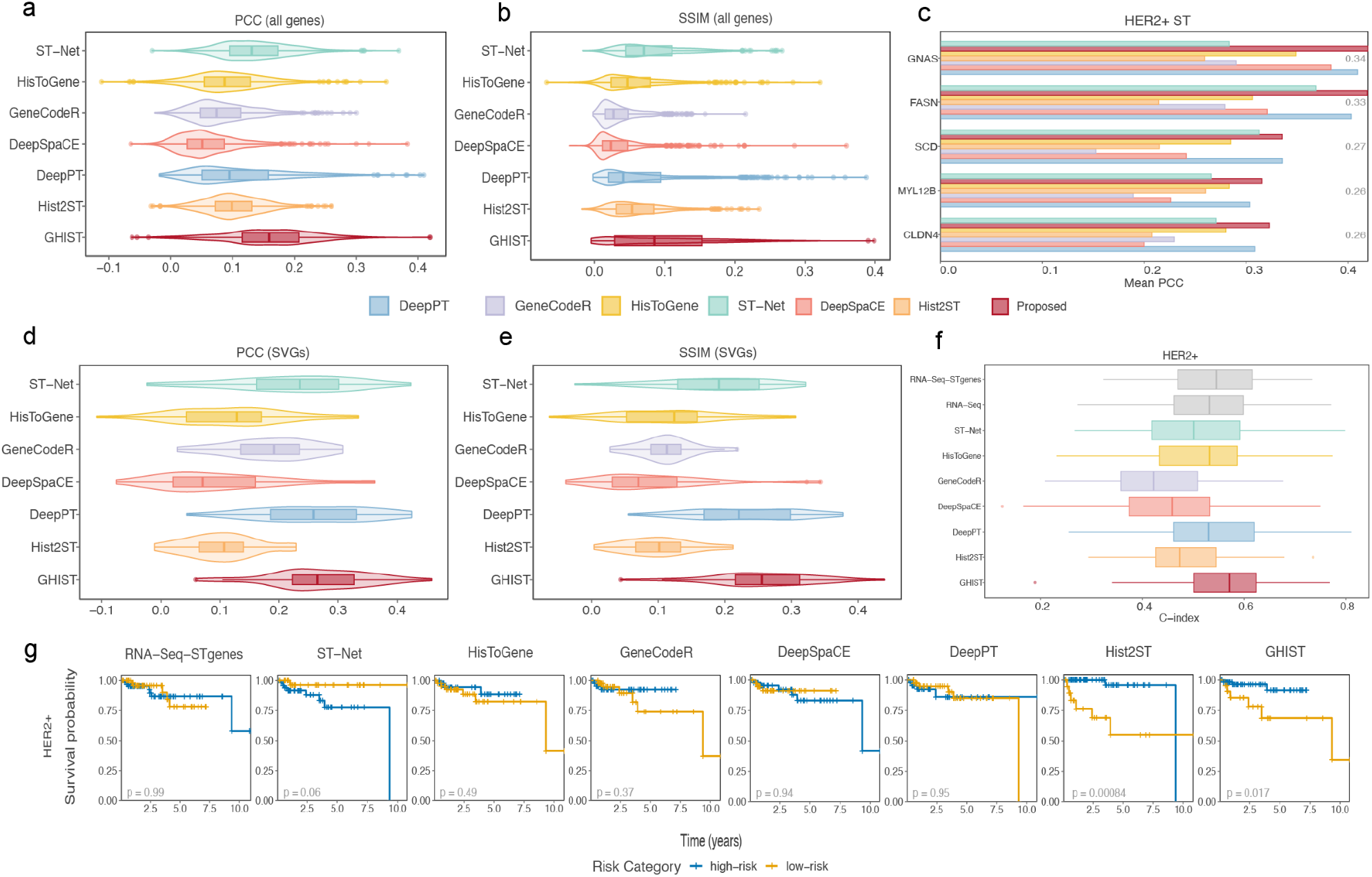
Comparison of spot-based gene prediction and survival analysis performance among state-of-the-art methods and GHIST using the HER2ST dataset. **(a)** Violin and boxplots of the average PCC and **(b)** SSIM between ground truth gene expression and predicted gene expression. Metrics measured from the test fold of a 4-fold CV, averaged over each gene across the dataset. **(c)** Top five correlated genes. **(d)** PCC and **(e)** SSIM violin and boxplots for each method for selected SVGs. **(f)** C-indices of multivariate cox regression models predicting survival of HER2+ subtype from TCGA-BRCA patients, using RNA-Seq bulk, RNA-Seq bulk using only genes present in HER2ST dataset, and the predicted pseudobulk from each method. C-indices were calculated from the test sets of a 3-fold CV with 100 repeats. **(g)** Cross-validated KM curves for patients split into high and low risk groups by the median risk prediction of the multivariate cox regression models for each method and HER2+ breast cancer subtypes. Each violin and boxplot in this figure ranges from the first to third quartile with the median as the vertical line. The lower whisker extends 1.5 times the interquartile range below the first quartile, while the upper whisker extends 1.5 times the interquartile range above the third quartile.

Moreover, we tested the translational potential of predicted SGE for downstream applications using selected TCGA-BRCA cancer individuals. Here we converted the predicted SGE to pseudobulk GE for each patient, and built multivariate Cox regression models from the pseudobulk GE to predict survival using a 3-fold cross validation with 100 repeats (**Figure 3f**). The corresponding RNA-Seq and a subset (RNA-Seq-STgene), comprising only genes present in the HER2ST dataset, were used as baselines for survival analysis. GHIST achieved the highest average C-index of 0.57, compared to 0.55 for RNA-Seq-STgene. Next, we binarised the risk model predictions of each patient and split them into high and low-risk groups based on the median risk prediction, then constructed Kaplan-Meier (KM) survival curves (**Figure 3g**). GHIST showed translational potential by demonstrating a significant difference in survival profiles between individuals from different risk categories (p = 0.017).

### GHIST is scalable and translatable when applied to TCGA data

To examine the GHIST model in practice, we also applied the model in its default single-cell expression setting directly to a TCGA-BRCA HER2+ cohort consisting of 92 patients (defined by patient metadata, details in Methods). The TCGA data is a multi-omics collection, containing histological images, bulk RNA-sequencing data, whole genome sequencing (WGS) data, and other information such as proteomics. We used the histological image as the input to obtain predicted expressions of 280 genes in individual cells. **Figure 4a** illustrates the predicted expressions for H&E patches from two individual samples. We show the UMAP of the cell type compositions for each individual in **Figure 4b** (top right panels). Next, we examine the location of the expression for two marker genes; *DSP* which is a marker of malignant cells, and *SFRP4*, which is more highly expressed in the adjacent tumour microenvironment (**Figure 4b** and **Supplementary Figure 7**). These observations are consistent with the general assumption that the tumour exists as a confined structure while other cell types such as fibroblasts and macrophages form the adjacent tumour microenvironment. **Figure 4c** shows the overall cell type proportion for all 92 individuals, consisting mostly of malignant cells, followed by stromal cells as expected. Two patients had minimal predicted malignant cells, and we found their H&E slides to have poor stain quality as assessed by a pathologist (**Supplementary Figure 8, Supplementary Table 1**, and Methods). Using GHIST, we generated spatially resolved single-cell expression of the TCGA data, thereby contributing a new modality to the TCGA multi-omics collection.

**Figure 4.**
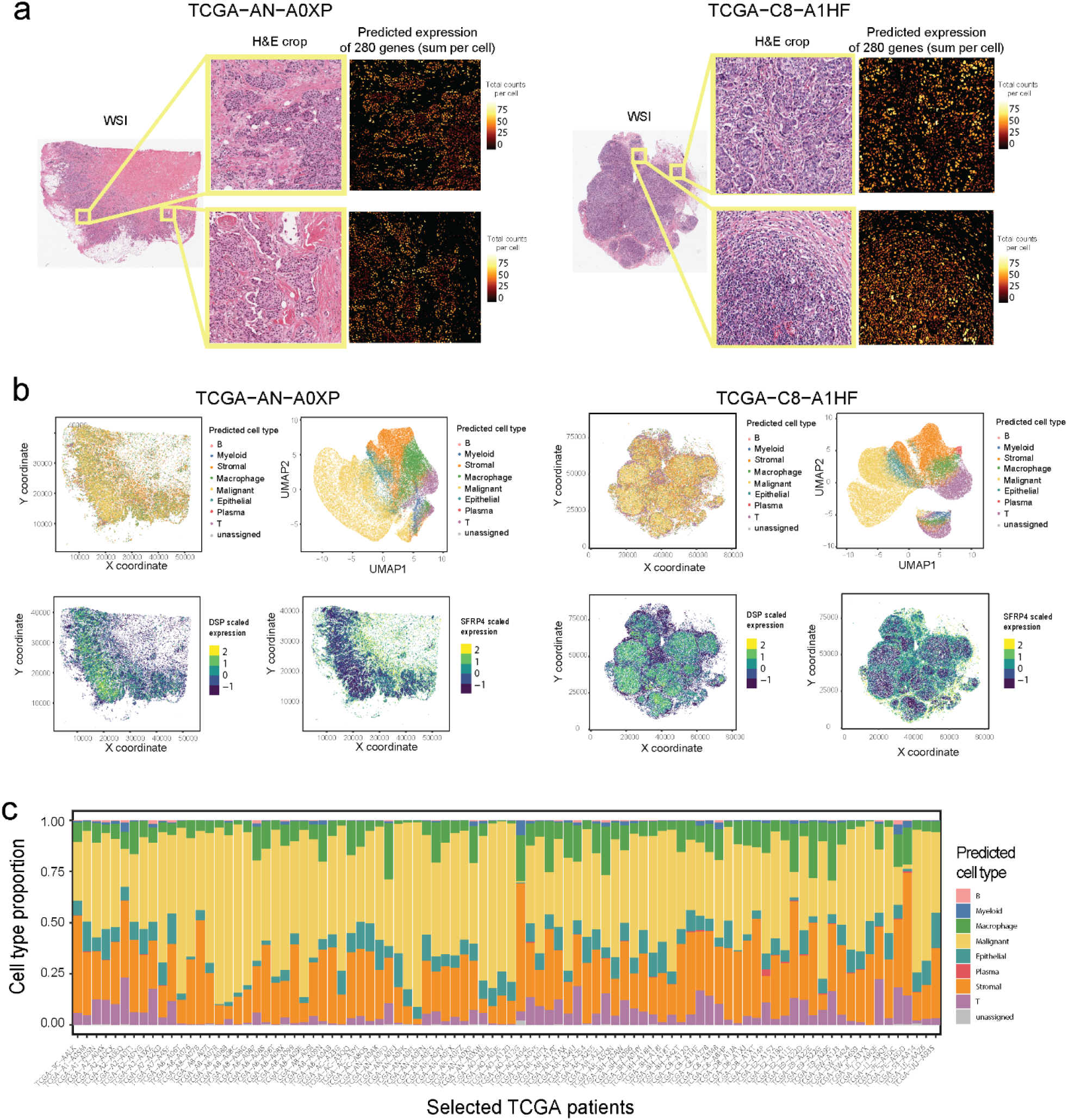
Application of GHIST to TCGA breast cancer H&E images. **(a)** Illustration of the total predicted gene expressions for each cell for a given H&E image taken from two TCGA-BRCA samples. **(b)** Cell type classifications using the predicted expressions, UMAP of the cell type compositions, and illustrations of the predicted expressions of *DSP* and *SFRP4* within cells in the two samples. **(c)** Predicted cell type proportions for the selected TCGA HER2+ patients.

### Potential of GHIST to create a *in silico* modality for multi-view and multi-omics analysis

We demonstrate the analytical potential of GHIST through multiple downstream analyses, such as survival analysis and differential state expression, using predicted single-cell data. Using the same selected subset of TCGA patients, we examined if the predicted gene expression is associated with survival outcomes. Firstly, we investigated the different types of features that can be extracted from the SGE predicted by GHIST. We revealed that the cell type information enabled by GHIST was able to differentiate high- and low-risk patients (**Figure 5a, Supplementary Figure 9**). While the cell type specific gene expression (chi-sq = 2.16) was marginally better in separating the risk groups compared with TCGA bulk RNA-seq (chi-sq = 1.9), we observed that L statistics (chi-sq = 6.34), which represents the spatial association of cell types, achieved a significant p-value of 0.012.

**Figure 5.**
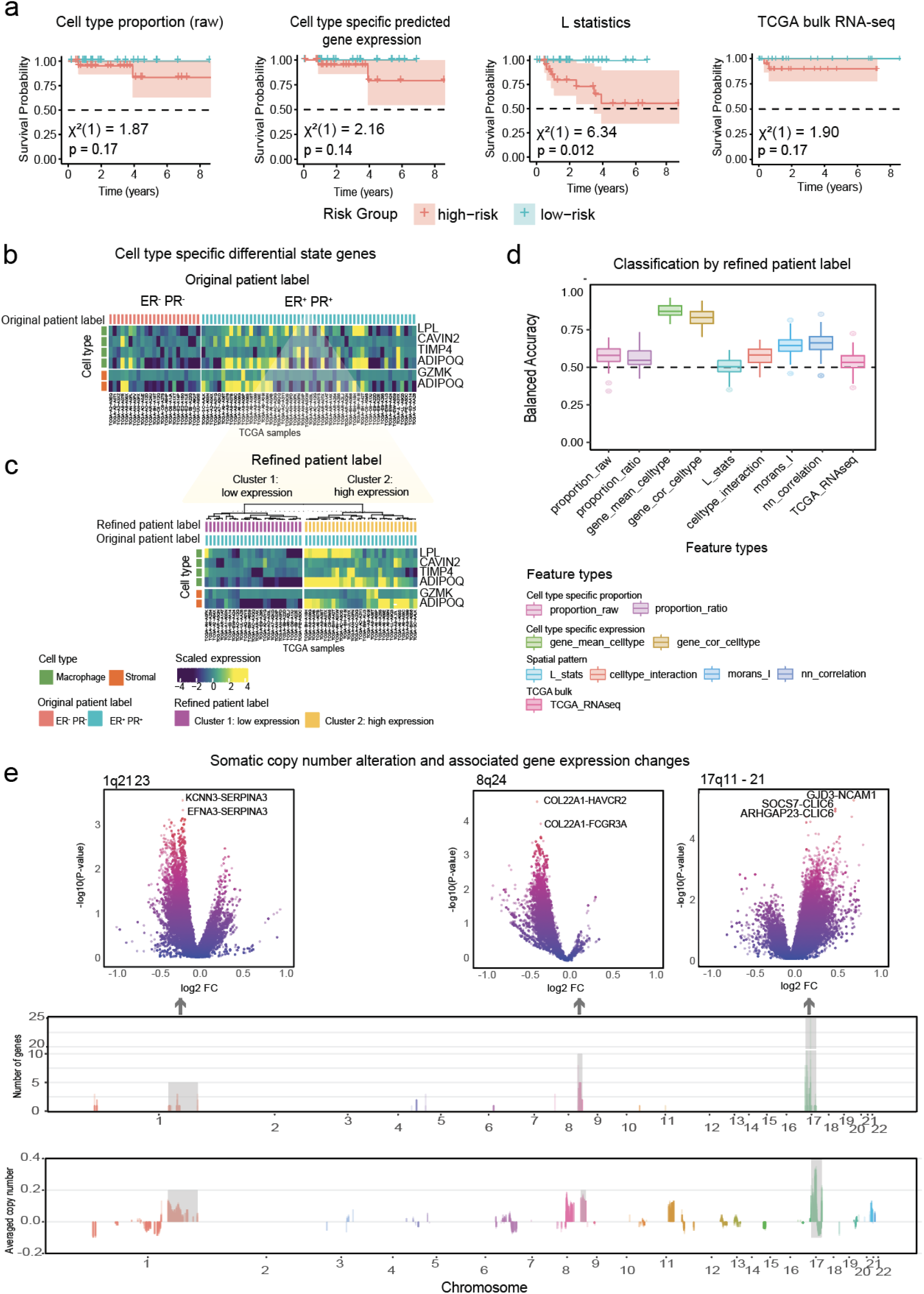
Potential of GHIST to create an *in silico* modality for multi-view analysis. **(a)** Cross-validated KM curves for patients split into high and low risk groups by the median risk predictions from cell type proportion, cell type specific predicted gene expression, L statistics of cell types, and RNA-seq data downloaded from TCGA. **(b)** Cell type specific differential state genes in macrophage and stromal cells between ER+/PR+ and ER-/PR- patients. **(c)** The ER+/PR+ group exhibited heterogeneity in expression of *LPL, CAVIN2, TIMP4, GZMK, and ADIPOQ*. Clustering method grouped them into two clusters. **(d)** Discrimination power of various feature types related to cell type proportion, cell type specific expression and spatial features between the two clusters. Each boxplot ranges from the first to third quartile with the median as the horizontal line. The lower whisker extends 1.5 times the interquartile range below the first quartile, while the upper whisker extends 1.5 times the interquartile range above the third quartile. **(e)** Differential spatial gene expression affected by copy number alteration. Volcano plots display 3 selected hotspots (1, 8, 17q) that affect a number of spatial expression patterns of genes (top). The number of differential patterning genes that were affected by each CNA (middle). Averaged copy number alteration profiles of 2,177 genes tested in the 91 patients are displayed (bottom).

Next, we examined disease subtype detection by comparing oestrogen receptor positive (ER+)/progesterone receptor positive (PR+) and oestrogen receptor negative (ER-)/progesterone receptor negative (PR-) patients, which are known to have distinct survival outcomes^24^ (**Figure 5b**).

Cell-type-specific analysis uncovered differential state genes specific to macrophage and stromal cells, such as *LPL, CAVIN2, TIMP4, ADIPOQ*, and *GZMK*. We further observed that these genes exhibited heterogeneity within the ER+/PR+ group, where one subgroup had lower expression similar to the ER-/PR- group, while the other subgroup had higher expression. We show that the high and low expression pattern is not a technical artefact through volcano plots, where there are a similar number of up-regulated and down-regulated genes between cluster 1 and cluster 2 for all cell types (**Supplementary Figure 10**). **Figure 5c** shows the results of unsupervised clustering on the ER+/PR+ individuals, demonstrating that cell-type-specific information enabled us to uncover patient heterogeneity within the ER+/PR+ population. In cluster 1, the higher expression of *LPL, CAVIN2*, and *TIMP4* within macrophage cells, and *ADIPOQ* in stromal cells, was associated with better survival (**Supplementary Figure 11a**). Conversely, lower expression was associated with poorer survival. This finding is consistent with the functions of these genes, where lower expression implies impaired metabolic and immune functions.

To further investigate the features whether there exists potential differential patterns between these two clusters, we employed a classification approach to assess the discriminatory potential of spatial patterns compared to cell type-specific expression patterns. We used scFeatures^25^ to generate a series of feature types and revealed that spatial metrics for differential spatial patterns, such as Moran’s I and nearest neighbour correlation (Moran’s I accuracy= 0.64 and nearest neighbour correlation accuracy = 0.66), also exhibited discrimination capacity (**Figure 5d)**. As expected, cell-type-specific gene expression (accuracy = 0.87) has a strong discrimination capacity as the clusters were initially defined by this measure. The spatial pattern-based feature type, nearest neighbour correlation, captures the composition and distribution of cell types in various local regions. Features that demonstrate differential patterning (as defined by nearest neighbour correlation) includes *TNFRSF17, CD19, KRT23, KRT7, TRAF4* (**Supplementary Figure 11b**), which all have a higher expression correlation with neighbouring cells in cluster 2 which correspond to better survival patients. These findings highlight the power of GHIST in uncovering additional spatial insights beyond the bulk gene expression data available in TCGA.

### Analysis between somatic CNA and NN correlation reveals that several genomic regions affect the spatial pattern of gene expression

Somatic copy number alteration (SCNA) has been revealed as a prognostic biomarker for breast cancer, and can often result in gene expression changes^26,27^. We extended the concept of differential expression analysis to define differential patterning (DP) genes as genes showing a difference in nearest neighbour correlation between samples of interest, where nearest neighbour correlation reflects the degree of spatial clustering or dispersion of different cell types. Here, we investigated the effect of gene gain/loss on spatial patterning of gene expression. First, we selected somatic copy number genes (with gain frequency >=5 and loss frequency >= 5 in 91 patients; one patient did not have CNA data) from the whole genome, which resulted in 2,177 genes in total. Next, for each of the SCNA genes, we performed a DP analysis to identify DP genes between samples showing gene gain versus samples showing gene loss for each SCNA based on the predicted expression of the 280 genes. We identified transacting hotspots associated with a large number of DP genes (p-value < 1.25 × 10^−3^, **Figure 5d, Supplementary Table 2**), termed DP hotspots including loci on chromosomes 1, 8, 17q. Our results recapitulated some well-known genomic regions that have been previously reported to be associated with cancer risk and gene expression. For example, the DP hotspots on chromosome 8 are around the known “8q24 gene desert” that has been associated with prostate, colon and breast cancer^28–31^. Other DP hotspots on chromosome 17 were located in 17q11-21, which happened to be the amplified region containing HER2. We observe that gain/loss *TOP2A* is associated with the spatial expression of 28 genes (**Figure 5e, Supplementary Table 3**). *TOP2A* is a cancer chemotherapy target^32^, and its copy number variation has been revealed as a predictive biomarker of chemotherapy of breast cancer^33,34^. Taken together, our results suggest that the predicted spatial gene expression pattern is a powerful new modality that enables previously unexplored questions relating copy number alterations and differential patterning.

## Discussion

Here we presented GHIST, a novel approach for predicting single-cell SGE from histology images. GHIST leverages multitask DL strategies to map H&E images (three channels, RGB) to a collection of cell-level spatially resolved transcriptomic images, by leveraging joint learning of cell nuclei morphology, cell type information, neighbourhood information, and gene expressions. GHIST is the first method to predict at single-cell resolution. We then demonstrated that GHIST can also be applied to spot-based resolution data and is able to outperform state-of-the-art spot-based approaches. We then showed that we can generate spatially resolved gene expression at single-cell level for breast cancer images in TCGA *in silico*, thereby contributing a new modality to enrich downstream analysis.

The application of GHIST to TCGA demonstrated a strong capacity for discrimination based on overall expression, cell type-specific expression, and varying spatial patterns. GHIST is able to achieve this even with a limited number of slides or samples. We believe this is possible because of the large amount of information observed across the whole tissue section during training and the large number of cells within each slide. In practice, the training data examined thousands of high-resolution patches from a slide containing about 80,000 cells, allowing the model to capture intricate details and subtle variations, and thus ensuring strong practical performance even when trained on very limited samples. While the number of slides is limited, we are effectively learning from a larger number of matched cells from H&E images and gene expression measurements. This contributes to the model’s ability to generalise across H&E images from different cohorts, leading to robust and reliable predictions in real-world applications. We anticipate an increase in model robustness with the availability of more training data.

The TCGA-BRCA samples have currently been assayed by 10 different experimental strategies, including genomics, transcriptomics and proteomics. Our spatially resolved gene expression measurements complement these existing modalities, providing a spatial omics modality across all of the samples in the collections. Further to this, H&E is a ubiquitous, routinely used histological technique applied in standard clinical practice to provide prognostic information that informs clinical management of nearly all cancers. As such, H&E is widely available with detailed clinical information in our health systems. GHIST can enrich these large collections of existing H&E data by enabling large-scale generation of spatially resolved gene expression profiles, facilitating comprehensive analysis and discoveries. As high quality training datasets continue to be generated, there is a clear opportunity to increase the availability of this additional spatial modality for other cancer types in the TCGA, and other databases and sample collections. This will enable researchers to investigate SGE patterns to formulate hypotheses about disease mechanisms and cell interactions, or alternatively, allow researchers to pre-examine potential results *in silico* before conducting costly and time-consuming experiments.

Many current H&E approaches are limited in their ability to identify a diverse range of cell types^35–37^. For instance, the dataset created by the Breast Cancer Segmentation Grant Challenge^38^ focuses on discriminating four key cell types: tumour, stroma, inflammatory, and necrosis. In contrast, we can predict double the number of cell types using our approach. This expanded capability provides a more detailed and nuanced understanding of the tissue microenvironment. It is important to note that the further expansion of identifiable cell types is tightly linked to the availability of data and the design of gene panels. Compared to scRNA-seq data, where thousands of genes are measured and many cell types are marked by the expression of dozens of genes, spatial transcriptomics gene panels with 300-500 non-specific genes may limit our capacity to discriminate cell types if marker genes are not present. For example, in our application to breast cancer, identifying myoepithelial cells, the absence of which distinguish *in situ* from invasive ductal carcinoma, is challenging because commonly stained markers such as *P63* are not available in the panel. This highlights the importance of customised gene panel design, a service offered by most vendors, and the power of newly emerging algorithms in this area (e.g., geneBasis^39^ and gpsFISH^40^). We believe that incorporating many of these considerations going forward by our community will greatly enhance the precision and utility of spatial gene expression predictions, allowing for more comprehensive and accurate tissue analysis.

The power of SGE prediction also lies in its potential to gain deeper insights into disease mechanisms through additional spatial information. This allows us to not only determine whether a particular gene is expressed but also where it is expressed and its relative patterning. Such capability enables us to explore whether specific gene expression patterns can further delineate breast cancer subtypes. Our results show that the spatial features including the L statistic (representing the spatial association of cell types), Moran’s I (representing the spatial patterning of gene expression), and nearest neighbour correlation (representing the correlation of genes with neighbouring cells), achieved high discriminating power in both survival and classification analysis. Such exploration would not be possible with the existing bulk RNA-sequencing data in TCGA. These findings underscore the power of GHIST in enabling researchers to examine histology images with a spatial resolution that was previously unattainable.

During application of GHIST to TCGA data, we have noticed quality issues with H&E slides, such as variable, under- or over-staining (**Supplementary Table 1**), which affect the predictions of a trained DL model. Thus there is a need to account for the quality of H&E slides prior to the application of our model, as interpretation of suboptimal slides should be avoided, as per current histopathology practice. It is known that the quality of H&E staining is dependent on artefacts that are attributed to multiple factors from formalin fixation to tissue processing, including thick and thin sectioning, processing temperatures and times, and batch effects from different tissue processors^7,41^. To increase clinical utility, there is potential to optimise various aspects of H&E protocols with the aim of increasing accuracy associated with SGE prediction. This includes both wet lab protocols, such as standardising staining techniques, optimal fixation and implementing rigorous QC measures, and computational approaches implementing advanced image pre-processing such as stain normalisation and leveraging foundation models trained on a large number of H&E images. Integrating these optimised protocols will lead to more reliable identification of spatial patterns in gene expression and deeper insights into disease mechanisms.

## Conclusion

Here, we introduced a multitask DL approach that enables the prediction of single-cell SGE from histology images by learning from H&E and SST images, and leveraging the relationships between multiple layers of cellular information. Our new GHIST framework outperformed current state-of-the-art SGE prediction methods based on spot-based technologies. Additionally, this framework offers a computational strategy to generate an *in silico* SGE platform with rich spatial information, enhancing existing databases. This facilitates the discovery of spatial associations between biomarkers, offering new insights into cellular biology and disease mechanisms.

## Online Methods

### Datasets and preprocessing

#### [A] Subcellular *in situ* spatial transcriptomics data

##### (i) BreastCancer1 and BreastCancer2

The 10x Xenium breast cancer datasets included in this study were downloaded from https://www.10xgenomics.com/products/xenium-in-situ/preview-dataset-human-breast (accessed 14 March 2024). BreastCancer1 was characterised as T2N1M0, Stage II-B, ER+/HER2+/PR−. BreastCancer2 was characterised as stage pT2 pN1a pMX, ER−/HER2+/PR−. The H&E image was aligned with its corresponding DAPI image for each dataset and visually verified. The overall pixel dimensions (*h × w*) of the H&E images were 25,778 × 35,416 for BreastCancer1, and 24,890 × 34,142 for BreastCancer2.

Nuclei were segmented from the H&E images using Hover-Net^37^ that was pre-trained on the CoNSeP dataset. The PyTorch weights of the segmentation-only model were downloaded from the official GitHub repository. The whole slide images (WSI) H&E were divided into non-overlapping patches that were 3,000 × 3,000 pixels in size (or smaller patch sizes along WSI border for remainder pixels), and Hover-Net was applied to the patches. Nuclei detected from two adjacent patches that overlap along the patch borders were merged together to a single nucleus, to avoid division of a nucleus into more than one segment.

BIDCell^42^ was applied to the transcripts and DAPI data to segment cells in the Xenium datasets and extract the gene expressions within whole cells. Low-quality transcripts for Xenium data with a phred-scaled quality value score below 20 were removed, as suggested by the vendor^43^. Irrelevant data including negative control transcripts, blanks, and antisense transcripts were filtered out. There were 313 unique genes for BreastCancer1 and 280 genes for BreastCancer2. For BIDCell, default parameter values from the exemplar file for Xenium and the provided single-cell reference file were used (both files were downloaded from the official BIDCell repository, version 1.0.3).

To find the corresponding gene expressions of cells detected from the subcellular SGE data and cells detected from H&E, the amount of overlap between nuclei from the two data types was computed. An H&E nucleus had a matching subcellular SGE nucleus if the amount of overlap between two corresponding nuclei was at least 50% of the area of the H&E nucleus. Cells with 0 total expression counts, cells classified as ‘unassigned’ by scClassify, and those with a nucleus smaller than 10 µ*m*^2^ were filtered out. This resulted in a total of 94k cells with single-cell expression for BreastCancer1, and 80k cells for BreastCancer2. The count matrix output for the segmented cells from BIDCell was used to classify the cell types.

##### (ii) LungAdenocarcinoma

The 10x Xenium lung adenocarcinoma dataset with multimodal cell segmentation was downloaded from https://www.10xgenomics.com/datasets/preview-data-ffpe-human-lung-cancer-with-xenium-multimodal-cell-segmentation-1-standard (accessed 21 February 2024). The H&E image was aligned with its corresponding DAPI image for each dataset and visually verified. The overall pixel dimensions (*h × w*) of the H&E image was 17,098 × 51,187. There were 377 unique genes in this dataset.

The same process used to segment nuclei from the H&E images for the breast cancer datasets was applied to the LungAdenocarcinoma dataset. We used the cell segmentation provided in the downloaded data bundle, that was achieved through the multimodal cell segmentation pipeline from 10x Genomics.

The same process was used to determine the corresponding cells detected in H&E and subcellular SGE data and perform filtering of cells for the LungAdenocarcinoma dataset as the breast cancer datasets. For LungAdenocarcinoma, overlap was calculated with cells in the SGE data, rather than nuclei. This resulted in a total of 89k H&E cells with single-cell expression for LungAdenocarcinoma. The count matrix provided in the downloaded data bundle was used to classify the cell types.

##### (iii) Melanoma

The 10x Xenium melanoma dataset was downloaded from https://www.10xgenomics.com/datasets/human-skin-preview-data-xenium-human-skin-gene-expression-panel-add-on-1-standard (accessed 4 April 2024). The H&E image was aligned with its corresponding DAPI image for each dataset and visually verified. The overall pixel dimensions (*h × w*) of the H&E image was 27,276 × 31,262. There were 382 unique genes in this dataset.

The same data processing steps as for the other Xenium datasets were applied to the Melanoma dataset. For this dataset, we used the gene expressions within nuclei. Post filtering, there were a total of 47k H&E cells with single-cell expression. The count matrix output for the segmented nuclei used to classify the cell types.

#### [B] scRNA-seq reference data sets used in cell type classification

For each data, we used scClassify^44^ version 1.12.0, a supervised single-cell annotation tool to annotate the cell types of each cell, that uses reference data with known cell types. The following scRNA-seq references are used for building the pre-trained model for cell annotations.

- *Single-cell breast reference* - a publicly available breast cancer dataset from 10x (https://www.10xgenomics.com/products/xenium-in-situ/preview-dataset-human-breast) as the reference to annotate the breast sample slides. The processing of the dataset is the same as detailed in our previous publication^42^.
- *Single-cell melanoma reference* - we used the pre-processed melanoma dataset obtained from Durante et al.^45^ as it contains comprehensive annotations. We used the annotation provided, and grouped “Class 1A Primary Tumor Cells”, “Class 2 PRAME+ Primary Tumor Cells”, “Class 1B PRAME+ Met Tumor Cells” and “Class 2 PRAME- Primary Tumor Cells” into “Tumor”; “M2 Macrophages”, “M1 Macrophages” and “Monocyte” into “Macrophages”; “CD8, T Effector Memory”, “CD8, T Resident Memory”, “Cytotoxic CD8”, “Naive T Cells” and “T Regulatory Cells” into “T Cells”.
- *Single-cell lung reference* - we used the pre-processed lung atlas data from the CZI CELLXGENE data portal (https://cellxgene.cziscience.com/collections/6f6d381a-7701-4781-935c-db10d30de293) as it is one of the largest collections of single-cell lung data. We used the level 2 annotation in the ann_level_2 metadata column. We selected the main cell types “blood vessels”, “airway epithelium”, “alveolar epithelium”, “fibroblast lineage”, “lymphatic EC”, “lymphoid”, “myeloid”, and “smooth muscle”. We grouped “airway epithelium” and “alveolar epithelium” into “epithelium”.

#### [C] HER2ST spatial transcriptomics dataset

The Human HER2+ breast tumour (HER2ST) dataset^23^ was measured using SRT to investigate SGE in HER2+ breast tumours, from which the original investigators discovered shared gene signatures for immune and tumour processes. The dataset consists of 36 samples of HER2+ breast tissue sections from 8 patients.

For the histology images, 224 × 224 pixel patches were extracted around each sequencing spot. For GHIST, the patches were centre-cropped at 112 × 112 pixels and resized to 256 × 256 pixels, so that the resolution is suitable for the nuclei segmentation model trained on NuCLS, and to be consistent with single-cell settings. For the SGE data of each tissue section, the top 1,000 HVGs for each section were considered and those genes that were expressed in less than 1,000 spots across all tissue sections were excluded. This resulted in 785 genes for the training of all models.

#### [D] NuCLS datasets

The NuCLS dataset^46^ was used to provide cell segmentation and cell type labels for the HER2ST dataset, to enable the training of GHIST to predict spot-level gene expressions. The NuCLS dataset was downloaded from https://github.com/PathologyDataScience/BCSS (accessed 18 February 2024), and contains annotations of over 220,000 nuclei from breast cancer H&E images from TCGA. The annotations include a mixture of region-level (rectangular) masks and nuclei-level (outlining nuclei boundary) masks for individual nuclei. The annotations were created through a crowdsourcing approach involving pathologists, pathology residents, and medical students. There were 1,744 H&E patches in the dataset. The “classes” level of annotations were used as cell type labels for nuclei. The classes include 6 cell types: “lymphocytes”, “macrophages”, “stromal cells”, “plasma cells”, “tumour cells”, and “other”.

The original nuclei masks provided in the NuCLS were unsuitable to be directly used for segmentation training, due to the presence of coarse rectangular nuclei masks. Hover-Net (pre-trained on the CoNSeP dataset, weights downloaded from the official GitHub repository) was applied to the NuCLS H&E patches to generate fine-grained nuclei-level masks (as pseudo ground truth masks). Due to inconsistencies in the sizes of H&E patches, four 270 × 270 patches were cropped from each image, from each corner of the original patch, and used as input to perform segmentation.

For every ground truth nucleus, centroids of pseudo nuclei within the area of the ground truth nucleus mask were found. The pseudo nucleus with the centroid closest by Euclidean distance to the centroid of the ground truth nucleus was deemed the corresponding nucleus. The ground truth cell type label was assigned to the matched pseudo nucleus mask and used to train Hover-Net from scratch to perform simultaneous nuclei segmentation and predict the cell type of the 6 types in the labels. Training was performed with a random training split of 90% of available patches, and default model hyperparameters were applied. The model was trained for a total of 100 epochs; only decoders for the first 50 epochs, and all layers for another 50 epochs, as per default settings. The model from the final epoch was applied to the HER2ST data to generate labels to train GHIST.

#### [E] TCGA-BRCA dataset

Data from The Cancer Genome Atlas Breast Cancer (TCGA-BRCA)^47^ were used to evaluate model generalisability. This dataset was chosen as the predictions on these H&E images could be evaluated through the matched gene expression, survival data, and other “omics”. TCGA-BRCA data was downloaded using the TCGAbiolinks package^48^ version 2.29.6.

RNA-Seq data was obtained by following query options: project = “TCGA-BRCA”, data.category = “Transcriptome Profiling”, data.type = “Gene Expression Quantification”, experimental.strategy = “RNA-Seq”, and workflow.type=“STAR - Counts”.

Histology images were obtained by the following query: project = “TCGA-BRCA”, data.category = “Biospecimen”, data.type = “Slide Image”, and experimental.strategy = “Diagnostic Slide”.

Clinical (metadata) with variables for defining breast cancer subtypes for samples were downloaded from the TCGAretriever package version 1.9.1.

Somatic copy number alteration was downloaded from UCSC Xena (Genomic Data Commons (GDC) TCGA Breast Cancer).

We only considered images that had one associated RNA-Seq entry and were either stage I, stage III, or stage IIA due to storage limitations. In our study, we focused on the HER2+ breast cancer subtype from TCGA-BRCA, as this was the subtype of the data (BreastCancer2 and HER2ST) used to train the model. We selected the HER2+ patients based on a positive entry in the “lab_proc_her2_neu_immunohistochemistry_receptor_status” metadata column ^49^.

Overall we considered 92 TCGA samples consisting of matched images, RNA-seq data, somatic copy number data, and clinical data.

H&E processing - Images were read using the openslide-python library, and non-overlapping 256 × 256 patches were then cropped from the highest resolution level of the WSIs. The patches were filtered out if the sum of the RGB values were greater (or more white) than a patch with RGB (220,220,220).

Nuclei were segmented from the patches using Hover-Net. We found that the stain colour affected the number of nuclei that the pre-trained model predicted, so the patches underwent stain normalisation. The torchstain package (version 1.3.0)^50^ was used to apply the Macenko normalisation method^51^ to each H&E patch. The reference was an unnormalised patch from a TCGA-BRCA H&E WSI for which Hover-Net performed well visually.

Macenko stain normalisation was also applied to the original TCGA H&E patches when input to GHIST. Here, the reference image was the whole H&E image from Xenium, downsampled such that 1 pixel = 10 μm.

### Spatially resolved single-cell Gene expression from HISTology (GHIST) Overview of GHIST

GHIST is a Deep Learning (DL) framework that considers the relationships between multiple layers of biological information to predict spatially resolved gene expressions within individual cells from visual information captured in H&E-stained histology images. GHIST uses two types of input data at inference: (i) H&E images, and (ii) optionally, average gene expression profiles of cell types from reference datasets, such as the Human Cell Atlas. During training, subcellular SGE data are used to quantify single-cell gene expressions for the paired H&E image to enable the framework to learn relationships between H&E images and gene expressions within cells. Once trained, the framework can infer single-cell SGE from H&E images without requiring SGE data.

A major **innovation** in developing GHIST is the harnessing of the complex relationships between the histological appearance of cells, gene expression within cells, cell type, and composition of neighbouring cell types. This multilayered approach enhances the capacity of the framework to better discern the spatial gene expression patterns reflected by H&E, thereby facilitating greater biological insights. These different types of information are integrated by considering them as prediction tasks in the multitask DL framework of GHIST. The primary task of GHIST is spatially resolved gene expression prediction within individual cells, and this is supported by three auxiliary helper tasks (morphology, cell type, and neighbourhood composition):

#### (A) Visual information that describe nuclei morphology and cell type

The visual appearance of tissues, cells, and nuclei reflects underlying biological information. GHIST extracts information regarding nuclei morphology, texture, and the appearance of the environment and neighbouring cells from H&E images. This is learned through a backbone network that performs joint nuclei segmentation and classification. Nuclei-level and patch-level features from the backbone are combined together. These features are fed into two auxiliary (cell type and neighbourhood composition) and primary prediction tasks downstream, so that these tasks receive a comprehensive embedding of information from H&E images.

The backbone is parameterised by a set of learnable parameters *θ* of a DL segmentation model. The backbone predicts the probability of cell type classes and background at each pixel. In this way, the model learns the relationships between pixels with morphology and cell type. We used the popular UNet 3+^52^ (without the deep supervision component) as the backbone of our framework to perform nuclei segmentation and classification. This backbone architecture may be swapped out for other segmentation architectures. The architecture comprised an encoding branch and decoding branch with five levels of feature scales, and incorporated full-scale skip connections that combined low-level details with high-level features across different scales (resolutions).

#### (B) Cell type information

The level of expression of genes in a cell is directly related to its cell type, as different cell types are known to be associated with distinct expression profiles and different marker genes. GHIST harnesses the relationship between gene expressions and cell type by carrying out cell type prediction as an auxiliary task in its multitask framework. This serves three key purposes: (i) incentivises the extraction of visual features that target cell type, which implicitly informs the gene expression profile of the cell; (ii) provides an alternative means of assessing and penalising the predicted expressions during training; and (iii) assists with the estimation of the composition of cell types within a local neighbourhood (see below for section *Neighbourhood characteristics*). As a result, the framework can also better extract meaningful features that improve the prediction of gene expressions.

To achieve point (ii), during training, the predicted gene expressions are used as input to another component in the framework to predict cell type. This allows the framework to learn single-cell expressions that reflect characteristic profiles that correspond to different cell types.

#### (C) Neighbourhood characteristics

Cells are known to exhibit particular patterns of spatial organisation in tumours, with different compositional patterns in local neighbourhoods^53,54^. Thus, we incorporate a module that learns to estimate the neighbourhood cell type composition of a given input patch as a further type of auxiliary output prediction. This provides two key benefits: (i) ensures that the combination of low-level features from individual cells is coherent with the local neighbourhood, and (ii) aids with discerning cell types that are visually challenging to distinguish in H&E images. As an example in breast cancer, B and T cell nuclei exhibit remarkably similar visual characteristics. B cells tend to be mistaken as T cells when the frequency of T cells is relatively dominant. These cell types often occur together in local neighbourhoods. We leverage this knowledge in GHIST by using the estimated neighbourhood composition to capture a more comprehensive view of the cellular environment in H&E images.

#### (D) Spatially resolved gene expression in individual cells

The integration of the three components above allows the main task, single-cell SGE prediction, to be facilitated by the different interrelated biological information. Once trained, GHIST predicts the expression of hundreds of genes for every nuclei detected in an H&E image, without requiring additional cell type, neighbourhood composition, or SRT information as inputs. GHIST can leverage averaged gene expression profiles of various cell types from single-cell references of the target tissue type as an optional input (ablation results in **Supplementary Figure 1**). The selection of the reference data is flexible, as GHIST does not require a matched single-cell reference for the same sample of interest, the cell types do not need to align with the cell types being predicted, multiple reference datasets may be used, and some reference datasets may contain different cell types.

#### [A] Extraction of visual information

The input to the model is a cropped patch from the H&E WSIs, *x* ∈ ℛ^*h*×*w*×3^, where *h* is the height of the patch, and *w* is the width of the patch, and 3 corresponds to the RGB colour channels. All the cells in an input patch are processed simultaneously during one forward pass, thereby allowing the model to flexibly support patches containing an arbitrary number of cells.

The backbone predicts an output of shape [*h, w, n*_*CT*_ + 1], where each element represents the probability of each cell type or background class at each pixel. During training, the backbone is trained carry out nuclei segmentation and cell type classification by minimising the loss function:

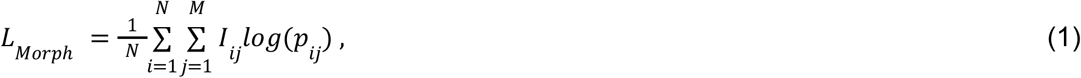

where *N* is the number of pixels, *M* is the number of classes (cell types, and background), *I*_*ij*_ is a binary indicator of whether class *j* is the correct classification for pixel *i* (1 if true, 0 otherwise), and *p*_*ij*_ is the predicted probability that pixel *i* belongs to class *j. M* is only relevant during training, and depends on the granularity of the cell types of interest. In our experiments, we found PCC of predicted SGE with ground truth to reduce for large *M* (see Discussion).

Nuclei-level features are extracted by multiplying nuclei masks for all valid nuclei in the input patch elementwise to the feature volumes produced by the first and last convolutional layers within the backbone architecture. The resulting features are concatenated along the feature dimension, then the features within each nuclei are summed and normalised by the size of the nucleus in pixels. This is done to counter the variability in nuclei sizes. This yields a unique feature vector with a dimension of 384. Furthermore, patch-level features are calculated by averaging the same two feature volumes across the patch. Each nucleus-level feature vector is concatenated with the patch-level feature vector of its corresponding patch. This is then processed through two fully connected layers with ReLU activation, to generate the resulting final feature vector that describes the morphology of each nucleus *x*_*nucleus*_ ∈ ℛ^1×256^. This describes nuclei-level and patch-level visual features that are relevant to nuclei morphology, cell type, and the local neighbourhood from H&E image.

#### [B] Leveraging cell type information

The cell type prediction module uses the extracted visual features of each nucleus, *x*_*nucleus*_, to predict the cell type of the nucleus. The module comprises a fully connected layer (256-dimensional intermediate features), followed by ReLU activation, and another fully connected layer that predicts the probability of each cell type. The argmax function is applied to obtain the predicted classes. This module is trained by minimising:

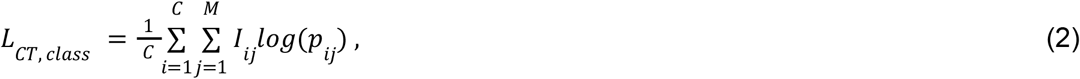

where *C* is the number of cells (i.e., nuclei), *M* is the number of classes (cell types), *I*_*ij*_ is a binary indicator of whether class *j* is the correct classification for cell *i* (1 if true, 0 otherwise), and *p*_*ij*_ is the predicted probability that cell *i* belongs to class *j*.

We introduce another module with the same composition of layers as the cell type prediction module, with three associated training losses, *L*_*CT,embed*_, *L*_*CT,logits*_, and *L*_*CT,expr*_ (defined below) that work in tandem. This module separately takes as input the predicted expressions and ground truth expressions, and predicts the corresponding cell type for each set of input expressions.

- *L*_*CT,expr*_ is the cross-entropy loss (Equation 2), computed exclusively when the inputs are the predicted expressions.
- *L*_*CT,embed*_ addresses the consistency of intermediate representations derived from the ground truth and predicted input expressions, and is written as:

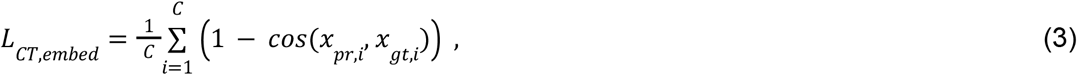

where *x*_*pr,i*_ is the embedding for the predicted (“*pr*”) expressions of cell *i*, and *x*_*gt,i*_ is the embedding for the ground truth (“*gt*”) expressions of the cell.

- *L*_*CT,logits*_ addresses the consistency between the predicted cell types derived from predicted and ground truth gene expressions for a given cell:

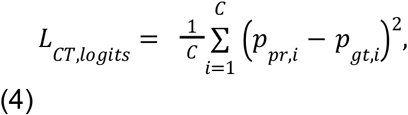

where *p*_*pr,i*_ is the output cell type logits given the predicted expressions of cell *i*, and *p*_*gt,i*_ is the output cell type logits given the ground truth expressions of the cell.

Taken together, predicted single-cell expressions are encouraged to resemble profiles that better correspond to cell types.

#### [C] Leveraging neighbourhood characteristics

GHIST estimates the composition of cell types in each input H&E patch. This module comprises two fully connected layers with ReLU activation, with the last layer outputting the same number of elements as the number of cell types in the training data. Each value represents the proportion of cells of a particular cell type in a patch. Softmax activation is used to ensure the compositions sum to 1 for each patch. The input to this module is the average of *x*_*fv, nucleus*_ for all nuclei in a patch.

This neighbourhood composition (NC) module is trained by minimising *L*_*NC,est*_ and *L*_*NC,pr*_, calculated for a given patch defined as:

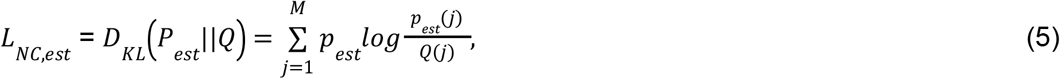

where *D*_*KL*_ is the Kullback-Leibler (KL) divergence between the cell type compositions estimated from the neighbourhood composition module *P*_*est*_, and the ground truth compositions *Q*, and *j* ranges over all the cell types. Similarly, for a given patch, the loss associated with neighbourhood composition module given the auxiliary predicted cell types:

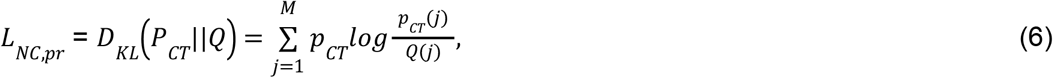

where *P*_*CT*_ is the composition calculated by summing the predicted cell types (see section *Leveraging cell type information*) and dividing by the total number of cells in a given patch.Thus, further consistency is imposed between the different layers of biological information within the model.

The estimated neighbourhood composition provides further utility in accounting for cell types that are difficult to discern from H&E images. For example, B cells and T cells have similar morphology, and thus the model becomes biased towards the dominant cell type (usually T cells), while ignoring the minority cell type (B cells). Based on this, during inference, we use the estimated compositions to recover missing cell types. For cell type *t* (e.g., immune cell types including B, myeloid, and T cells):

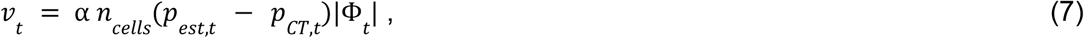

where *n*_*cells*_ is the number of cells in the patch, *Φ_y_* ∼ *N*_*t*_ *(p*_*est,t*_, 1) is sampled from a normal distribution, where *n*_*I*_ is the number of B, myeloid, and T in the patch from the auxiliary cell type predictions, and *α* is an adjustable hyperparameter to allow different recovery rates. We set *α* to 2 in our experiments for B, myeloid, and T cell types, and T cells initially predicted with high confidence (probability above 0.6) were masked from this mechanism. These values were selected empirically, according to performance on the validation sets. Next, the logits of the initial predictions *y’*_*CT,tc*_ for the cells are revised as follows:

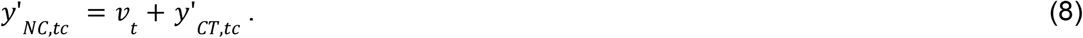

The final cell type is found by applying argmax across the cell types being refined, while not affecting other cell types. To ensure consistency, the final expressions are taken from the corresponding cell type-specific prediction module (see section on *Gene expression prediction*). Disease-specific knowledge may also be incorporated into this mechanism (Equations 7 and 8), e.g., decreasing the relative proportion of epithelial to malignant cells in invasive breast cancer. We only demonstrate this mechanism on the breast cancer datasets; it was not applied for the dataset Melanoma or LungAdenocarcinoma.

#### [D] Spatially resolved gene expression prediction for individual cells

GHIST predicts the expression value of every gene in each cell in the H&E image, 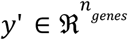. The number of genes predicted varies according to the size of the gene panel in the available SST data. The datasets used in our study had panel sizes ranging from 280 to 382 genes.

The input to this module is the embedding vector *x*_*nucleus*_ ∈ ℛ^256^ for all nuclei in a patch (with dim = 256), and if used, a set of averaged gene expression profiles of various cell types from single-cell references of the target tissue type 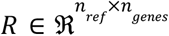, if this input is used. The reference data does not need to be matched for the same sample of interest, and *n*_*ref*_ can represent a flexible number and categories of cell types from a single or multiple reference datasets. *R* is kept the same during training and inference.

GHIST carries out a linear regression to predict *y’* by first predicting 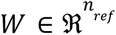:

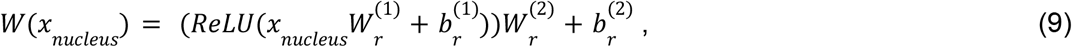

followed by applying softmax along the *n* dimension, where 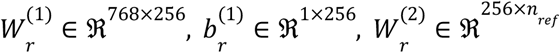, and 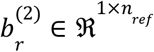. The *S*_*j*_ is the expression value of gene *j* calculated by the weighted sum of references for each cell written as:

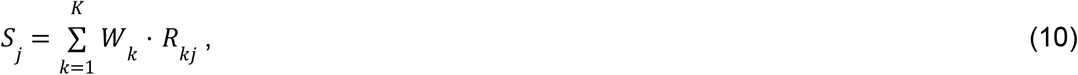

In the setting without using single-cell reference as an input:

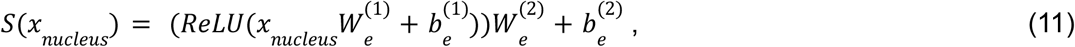

where 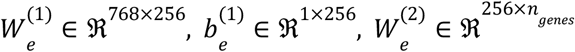, and 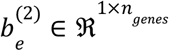. Next, information about the spatial neighbourhood is incorporated using the estimated cell type neighbourhood composition 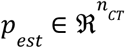 of a patch. We use multi-head cross attention (with 8 heads), which are fundamental components in Transformers. For each cell *p* forms the query, while *S* forms the key and value.

A linear layer is applied to the output of the multi-head cross attention module, resulting in 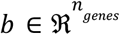. Finally, the predicted gene expressions of each cell are computed:

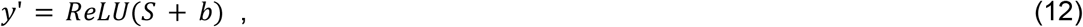

In the setting without using single-cell reference as an input:

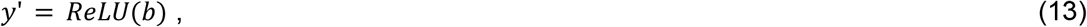

For the spot-based prediction mode, *y’* for all nuclei in a patch are summed together (for weakly-supervised learning):

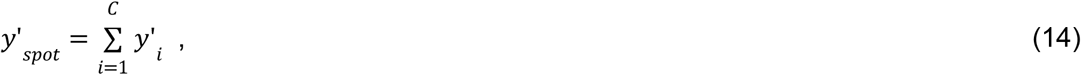

The training loss associated with gene expressions is:

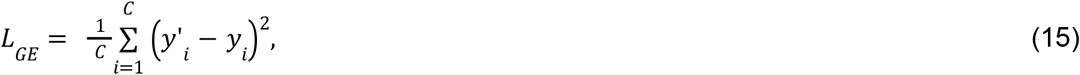

where *y*_*i*_ is the gene expressions for cell *i* obtained from SST. When the estimated neighbourhood compositions are used to recover missing cell types, a copy of the components from Equations 9 to 13 is made for these specific cell types. In this case the loss *L*_*GE,adj*_ is calculated only where these cell types are present.

#### [E] Model Training

The model was trained by minimising the **total loss** representing sum of all the losses over *N* training patches:

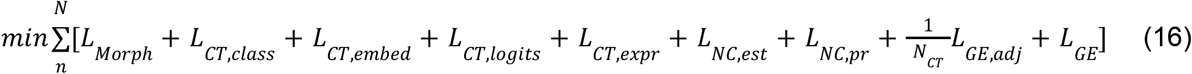

Due to the relatively small value of *L*_*CT,embed*_ observed during training, it was empirically scaled by a factor of 100. *L*_*GE,adj*_ was inversely scaled by *N*_*CT*_, the number of cell types in the auxiliary prediction. There were no further weightings for the other losses, to improve generalisability and ensure that our losses were not fine-tuned to any particular datasets.

### Practical implementation of GHIST

Cross-validation (CV) was carried out for the Xenium and HER2ST datasets. The training set was used to train parameters of the model. The validation set was used to evaluate models during training to allow parameters to be tuned. Results presented in the paper were calculated using the predictions using the test set for each of the four folds of the CV. The same CV splits were used for each model to ensure fair comparison.

5-fold CV was carried out for each Xenium dataset. The whole H&E image in each dataset was horizontally sectioned into 5 non-overlapping portions, with each portion serving as the testing data for a fold. The remaining portions formed the training and validation sets, where the validation set was a section that was a tenth of the whole H&E in height. The training portion of a fold was divided into non-overlapping 256 × 256 patches. During inference, patches were divided such that there was a 30 pixel overlap (approximately the width of the largest nuclei in the data). The final prediction for each distinct nucleus was based on the input image containing the largest area of that particular nucleus.

For the HER2ST dataset, the SGE prediction was evaluated using 4-fold CV. The dataset was split into four folds, with samples from the same patient belonging in the same fold. In each iteration of the CV, two folds were used as a training set, one fold used as a validation set and the remaining fold used as a test set. The best model within each fold was then chosen based on the epoch that produced the best PCC in the validation set.

For GHIST, batch size was set to 8 during training and inference. The model is flexible regarding the number of nuclei present in the patch, and outputs predicted gene expressions as *n*_*cells, total*_ × *n*_*genes*_. The model was trained end-to-end from scratch for 50 epochs. The saved checkpoints (per epoch) were applied to the validation sets, and selection of the top performing checkpoint was based on the best average of the rank in F1 score (of the auxiliary cell type classification prediction component) and MSE (of the predicted single-cell expressions). The top performing checkpoint was then applied to the test set of the fold.

Weights of the convolutional layers in GHIST were initialised using He et al.’s method^55^. Input images were standardised using the mean and standard deviation of the training patches, and gene expressions were log-normalised. We employed standard on-the-fly image data augmentation by randomly applying a flip (horizontal or vertical), rotation (of 90, 180, or 270 degrees) in the (x,y) plane. The order of training samples was randomised prior to training. We employed the AdamW optimiser^56^ to minimise the sum of all losses at a fixed learning rate of 0.001, with a first moment estimate of 0.9, second moment estimate of 0.999, and weight decay of 0.0001.

We ran GHIST on a Linux system with a 24GB NVIDIA GeForce RTX 4090 GPU, Intel(R) Core(TM) i9-13900F CPU @ 5.60GHz with 32 threads, and 32GB RAM. GHIST was implemented in Python using PyTorch.

#### Application to TCGA-BRCA

Disease-specific knowledge was incorporated during inference on the TCGA images, due to the disparity between the sample that the model was trained on (which comprised normal ducts, ductal carcinoma in situ (DCIS), and invasive carcinoma), and those of the TCGA cohort (all invasive subtypes). Without fine-tuning, we set *α* to an arbitrary high value (10,000) and considering *p*_*CT*_ as 0 for malignant cell types (Equations 7 and 8). During inference on TCGA images, ensemble averaging was carried out using the model checkpoints from three folds that were trained on BreastCancer2.

### Performance evaluation

#### Evaluation study 1: Ablation study

We performed an ablation study to determine the contributions from each of the types of biological information in GHIST. We used BreastCancer2 for these experiments. We evaluated GHIST without neighbourhood composition prediction, thereby impeding the ability to refine predicted gene expressions for cells based on estimated neighbourhood characteristics. We also investigated the effects of performing auxiliary cell type prediction, by removing the ability of the model to leverage the relationships between gene expressions and cell type labels. Furthermore, instead of performing a linear regression of average cell type profiles, we investigated the setting where gene expressions are directly predicted as a straightforward regression.

We performed cell type classification using scClassify using the breast cancer reference data, followed by comparison of cell type distribution and cell type proportion with the ground truth data and the correlation of SVGs.

### Evaluation study 2: Comparison study with other spot-based methods

We describe the settings for other spot-based methods below, and more details may be found in our benchmarking work^15^:

#### [A] Settings used for other methods

- ST-Net^10^ - ST-Net was implemented in Python and PyTorch using hyperparameter values as outlined in the original paper. The DenseNet-121 backbone used was pre-trained on ImageNet. ST-Net was trained using the MSE loss function, computed between predicted and ground truth expression levels.
- HisToGene^9^ - The HisToGene model was implemented using PyTorch with the following hyper-parameters: a learning rate of 10e-5, the number of training epochs of 100, a drop-out ratio of 0.1, 8 Multi-Head Attention layers and 16 attention heads. The final model used was the model after all epochs, as per the HisToGene pipeline.
- GeneCodeR^13^ - GeneCodeR was the only method that did not use deep learning for its model. The following hyperparameters were used for the coordinate descent updating algorithm: the dimensions of the encoding space for the samples (*i_dim*) of k=500, the dimensions of the encoding space for the features (*j_dim*) of c=500, initialisation for the alpha_sample = “rnorm”, initialisation for the beta_sample = “irlba”, max iterations of 20, tolerance of 5, learning rate of 0.1, and batch size of 500.
- DeepSpaCE^12^ - The DeepSpaCE model was then implemented using the VGG16 model as the backbone with the following hyperparameters: maximum number of training epochs of 50 (with training stopping before 50 if there is no improvement within 5 epochs), a learning rate of 10e-4 and a weight decay of 10e-4.
- DeepPT^8^ - DeepPT was implemented in Python and PyTorch using hyperparameter values as outlined in the original paper. The ResNet-50 backbone used was pre-trained on ImageNet. DeepPT was trained using the MSE loss function, computed between predicted and ground truth expression levels.
- Hist2ST^11^ - The model was implemented using Python and PyTorch and ran with default parameters stated in the original paper. Some hyperparameter changes were made for benchmarking including: increased the image patch size from 112×112 to 224×224 and reduced the number of input and output channels in the depthwise and pointwise convolutional blocks from 32 to 16 due to limited computational power. Image embedding dimension was increased from 1024 to 2048 as the original code of setting embedding size was hard coded based on the image patch size and the number of channels. The model was trained using a sum of MSE loss of predicted and ground truth expression levels, ZINB and Distillation mechanisms.

We used code from the official GitHub repositories for each method unless otherwise stated.

#### [B] Evaluation measures

Concordance between predicted and measured gene expression - We followed the strategies described in our benchmarking work^15^ to assess the performance of GHIST on HER2ST data. Briefly, the evaluation methods used include PCC and SSIM. These metrics were first calculated at an image level (e.g. correlation was measured for each gene per image), and then averaged over each patient, then averaged over each gene.

HVGs and SVGs were deduced from the ground truth SGE from the HER2ST dataset. The *modelGeneVar* function, followed by the *getTopHVGs* function with prop=0.1 from the *scran* R package^57^ version 1.32.0 were used to obtain the HVGs in each ST dataset. For SVGs selection, genes with adjusted p<0.05 were identified as SVGs in each image sample in each dataset using the SPARK-X package in R^58^ (version a8b4bf2). For HER2+, only the SVGs that appear in more than 30 out of 36 samples were selected and correlation was calculated based on top 20 genes ranked by adjusted p-value.

Assessment of the translational potential with cross-validated (CV) C-index and KM curves - We first selected the top 20 genes associated with the highest C-indices from the univariate Cox models built on the predicted SGEs for each gene. Next, a multivariate Cox regression was then built using these genes. Each model was evaluated by C-index and a logrank p-value based on the predicted SGE for each of the spot-based methods and GHIST, where C-index was calculated by comparing the ground truth event times and events with model predictions for each patient within a test set.

While many existing methods^59^ perform C-indices calculation similar to a resubstitution model, we have used a three-fold CV with 100 repeats. This meant that patients were randomly split into three subsets. Each subset was used as a test set of patients while the rest of the patients were used to train models. Through repeating this process 100 times, we measured the C-indices for each test set to obtain an average C-index across 300 values. We also obtained the average prediction for each patient over all repetitions under CV which was then used to categorise patients into high and low-risk. KM curves were then constructed for each of the high-risk and low-risk groups. The log-rank test was then used to obtain a p-value to quantify the difference between survival of the two groups.

### Evaluation study 3: Performance evaluation at single cell resolution

Visual assessment - We selected cell type specific marker genes using the Findmarker function in Seurat to identify markers for each cell type. We visualised the predicted gene expression and the measured gene expression of the selected top marker genes.

Concordance between predicted and measured cell types proportion - We predicted the cell types in the TCGA data using the single-cell breast reference as aforementioned. We then visually compared the cell types between predicted and measured using barplots and scatterplots.

Correlation between predicted and measured gene expression - We treated the measured gene expression from the SST data as the ground truth. We computed the Pearson correlation coefficient (PCC) of a collection of genes between the predicted gene expression from H&E and the measured (ground truth) gene expression from SST. We calculated the PCC based on (1) top 20 SVGs, (2) top 50 SVGs, (3) top 20 HVGs, (4) top 50 HVGs, (5) top 20 non-SVGs, (6) top 50 non-SVGs, (7) top 20 non-HVGs and (8) top 50 non-HVGs. We defined the top SVGs and HVGs using the default ordering of the genes from the SVGs and HVGs calculation. The non-SVGs and non-HVGs are defined as the genes ranked in the bottom of the SVGs and HVGs calculation. For example, bottom 20 non-HVGs are defined to be the genes ranked in the bottom 20 of the HVGs calculation.

The SVGs were calculated using Giotto rank^60^ version 1.1.2, one of the top performing methods based on a recent benchmarking paper^61^. The HVGs were calculated based on the function FindVariableFeatures from the Seurat R package^62^ version 5.0.3.

Assessment of the translational potential of spatial features - We examined whether the prediction expression contains information that can be used to stratify patients into distinct survival groups. We used death as the survival event outcome, days to death as the survival times, and days to last follow up when there was no death event. We considered each of the feature types as described in the *Multi-view feature generation* section and **Supplementary Table 4** and examined the discriminatory power of each feature type by estimating the high-risk and low-risk individuals based on a given set of features.

For a given feature type, we selected the top 1% of the features, or the top 5 features, whichever number is greater, from each feature type separately (**Supplementary Table 4**) and used them to fit a Cox Proportional Hazard (CoxPH) model (function crossValidate in the ClassifyR^63^ R package version 3.9.1). Next, for each of the feature types described in **Supplementary Table 4**, we used 5,000 repeated three-fold cross-validation strategies to identify high-risk and low-risk individuals. In each iteration (one repeat), two folds of the patients were used to train the CoxPH model based on the selected features and individual risk was calculated on the remaining testing fold. This was repeated 5,000 times, resulting in a vector of a “risk score” for each patient. For each patient, this resulted in a vector of 5,000 risks estimated when the individual was inside the “test set” for each iteration. The final risk score was calculated as the median of this vector of risk estimates.

Using the final risk score, patients with a risk in the 0 to 25% quantile are defined as the “low risk group,” while individuals in the 75% to 100% quantile are defined as the “high risk group.” A Kaplan-Meier (KM) plot was then constructed, and a log-rank test was performed. The chi-square statistic and p-value associated with the log-rank test were calculated to measure the difference in survival between the two groups. We used the chi-square statistics associated with each feature type to assess the discrimination capacity of each feature type. To ensure a fair comparison with TCGA, we first subset the TCGA bulk RNA-seq to the same set of genes as in the H&E gene panel.

### Downstream statistical and computational biology analysis

To examine the downstream application of GHIST, we applied GHIST trained on the dataset BreastCancer2 to predict the expression on 92 TCGA patients (detailed in previous sections).

### Multi-view feature generation

We generated multiple types of features based on the prediction gene expression to enable more comprehensive exploration of the TCGA patients. Using the R package scFeatures^25^ version 1.4.0, we generated a total of eight feature types from three distinct categories: single-cell based cell type proportion, single-cell based cell type-specific expression, and spatial pattern. Each feature type contained tens to hundreds of features. **Supplementary Table 4** describes each feature type in detail.

### Differential state analysis between conditions (ER-/PR- vs ER+/PR+)

While we now have a computationally generated multi-sample, multi-group spatially resolved scRNA-seq TCGA-BRAC data, we first focused on identifying markers for differential cell states for each cell type between individuals with oestrogen receptor negative (ER-)/progesterone receptor negative (PR-) vs. oestrogen receptor positive (ER+)/progesterone receptor positive (PR+). We modelled this using a negative binomial generalised linear model implemented in the function pbDS in the R package muscat (version 1.18.0). Significant differential cell state markers were defined as genes with log fold change > 1 and p-value < 0.05. We visualised the expression of these cell type specific differential state genes as a heatmap.

### Unsupervised learning to detect novel sub-cohort in ER+/PR+

Within the ER+/PR+ cohort, we performed k-means clustering based on all cell type specific differential state genes identified in the previous section to cluster into two groups. We used PCC as the similarity metric and used the default parameters for the rest. This was performed using the Heatmap function implemented in the ComplexHeatmap^64^ R package version 2.20.0.

The characteristics of the two resulting clusters were determined in three different ways:

(a) Survival analysis - We calculated the KM curve for survival for cluster 1 and cluster 2 using the function survfit in R package survival version 3.6-4.

(b) DE analysis - We generated a large collection of features for each individual in the TCGA data using scFeatures (see *Multiview feature generation* section). Next, we identified features with a differential based on a moderated t-test using the lmFit function implemented in the package limma (version 3.60.2). We visualised these results using volcano plots.

(c) Classification analysis - We further performed classification to examine the separability of the two subgroups based on various feature types. To avoid the potential issue of “double dipping”, we removed all differential state genes from the features in each feature type, as well as genes with a correlation above 0.7 with any of the differential state genes. We assessed the performance based on balanced accuracy using 100 repeated three-fold cross-validation implemented in the crossValidate function in the R package ClassifyR. The crossValidate has an in-built feature selection using t-tests to select features with differential “measurement” between cluster 1 and cluster 2. We used a support vector machine (SVM) as the classification model backbone. For visualisation of most important features contributing to the classification, we selected top features based on frequency of inclusion, and visualised the values of these features in cluster 1 and cluster 2 samples using boxplots.

### Trans-acting associations between copy number aberration and nearest neighbourhood correlation

Focal copy number alteration (CNA) was estimated using the Genomic Data Common (GDC) pipeline with the GISTIC method implemented and then thresholded to -1, 0, 1, representing deletion, normal copy, or amplification. Here we selected 91 TCGA HER2+ patients that had copy number alteration results to perform downstream analysis, as one patient (TCGA-AR-A1AT) did not have CNA data. CNAs on chromosome X were removed. We first selected all genes that have both deletion and amplification with a frequency >= 5, which resulted in 2,177 CNA genes. Next, for a given CNA_gene, we compared between two groups of individuals, patients showing CNA_gene loss (−1) and CNA_gene gain (+1). For these two groups, we identified differential patterning (DP) genes where spatial patterning was measured by NN correlation. This DP analysis was repeated for all 2,177 CNA_genes a matrix of p-values representing the moderated t-tests for 2,177 gene CNAs being tested against 280 genes’ NN correlation (**Supplementary Table 2**). Due to the correlation between genes in the same genomic region, we did Bonferroni correction for 800 unique genes among the 2,177 genes. Genes with p-value < 1.25 × 1*0*^−3^ (1/800) were considered as DP genes, representing genes that significantly affect spatial expression patterns for a given CNA_gene. The number of DP genes are tabulated for each CNA gene and CNA clusters are identified as DP hotspots. Three hotspots regions were selected for highlighting differential NN correlation between gain and loss copy number by calculating -log2(fold change).

## Competing interests

The authors declare that there are no competing interests.

## Data availability

All datasets used in this study are publicly available and were downloaded from the following links. 10x Genomics Xenium breast cancer samples 1 and 2, and single-cell data: https://www.10xgenomics.com/products/xenium-in-situ/preview-dataset-human-breast. 10x Genomics Xenium lung adenocarcinoma:

https://www.10xgenomics.com/datasets/preview-data-ffpe-human-lung-cancer-with-xenium-multimodal-cell-segmentation-1-standard. 10x Genomics Xenium melanoma: https://www.10xgenomics.com/datasets/human-skin-preview-data-xenium-human-skin-gene-expression-panel-add-on-1-standard. Melanoma single-cell reference data was downloaded from dbGaP under accession code phs001861.v1.p1

(http://www.ncbi.nlm.nih.gov/projects/gap/cgi-bin/study.cgi?study_id=phs001861.v1.p1). Lung atlas data from CZI CELLXGENE data portal:

https://cellxgene.cziscience.com/collections/6f6d381a-7701-4781-935c-db10d30de293. HER2ST dataset: https://github.com/almaan/her2st. NuCLS dataset: https://github.com/PathologyDataScience/BCSS. TCGA-BRCA: https://portal.gdc.cancer.gov/projects/TCGA-BRCA. TCGA-BRCA clinical data: https://www.cbioportal.org/study/clinicalData?id=brca_tcga. Somatic copy number alteration for study name GDC TCGA Breast Cancer (BRCA) was downloaded from UCSC Xena https://xena.ucsc.edu/.

## Code availability

We provide our code for GHIST in https://github.com/SydneyBioX/GHIST.

## Acknowledgments

The authors thank all their colleagues, particularly at The University of Sydney, Sydney Precision Data Science and Charles Perkins Centre for their support and intellectual engagement. Special thanks to Matthew Shu, Agus Salim, and Hao Wang and for their contribution to discussions, and to Nick Robertson for testing the GHIST code.

## Funding

This work is supported by Judith and David Coffey Life Lab (CPC) to YC and JY; the AIR@innoHK programme of the Innovation and Technology Commission of Hong Kong to JY, JK, EP, YC, XF; NHMRC Investigator Grant APP2017023 to JY and CW; Chan Zuckerberg Initiative Single Cell Biology Data Insights grant (DI2-0000000197) to YC and JY; ARC Discovery Project DP200103748 to JK, and the USyd–UofG Ignition Grant to JK, JY, and XF.

The funding source had no role in the study design; in the collection, analysis, and interpretation of data, in the writing of the manuscript, or in the decision to submit the manuscript for publication.

## Author contribution

JY conceived and led the study with design input from XF. XF led the method development and implementation of the framework, with input from YC, JK and JY. YC and XF co-led the data assessment with input from JY, NP, DG, and EP. YC led downstream analysis with input from JY, XF, EP, DG and BB. BB led the multi-omics analysis with input from YC, XF, JY, DG and EP. CW led the spot-based evaluation. NP assessed the QC of all H&E slides and with DG, guided the refinement of the model and interpretation. All authors contributed to the writing, editing, and approval of the manuscript.

## Supplementary Figures

**Supplementary Figure 1.**
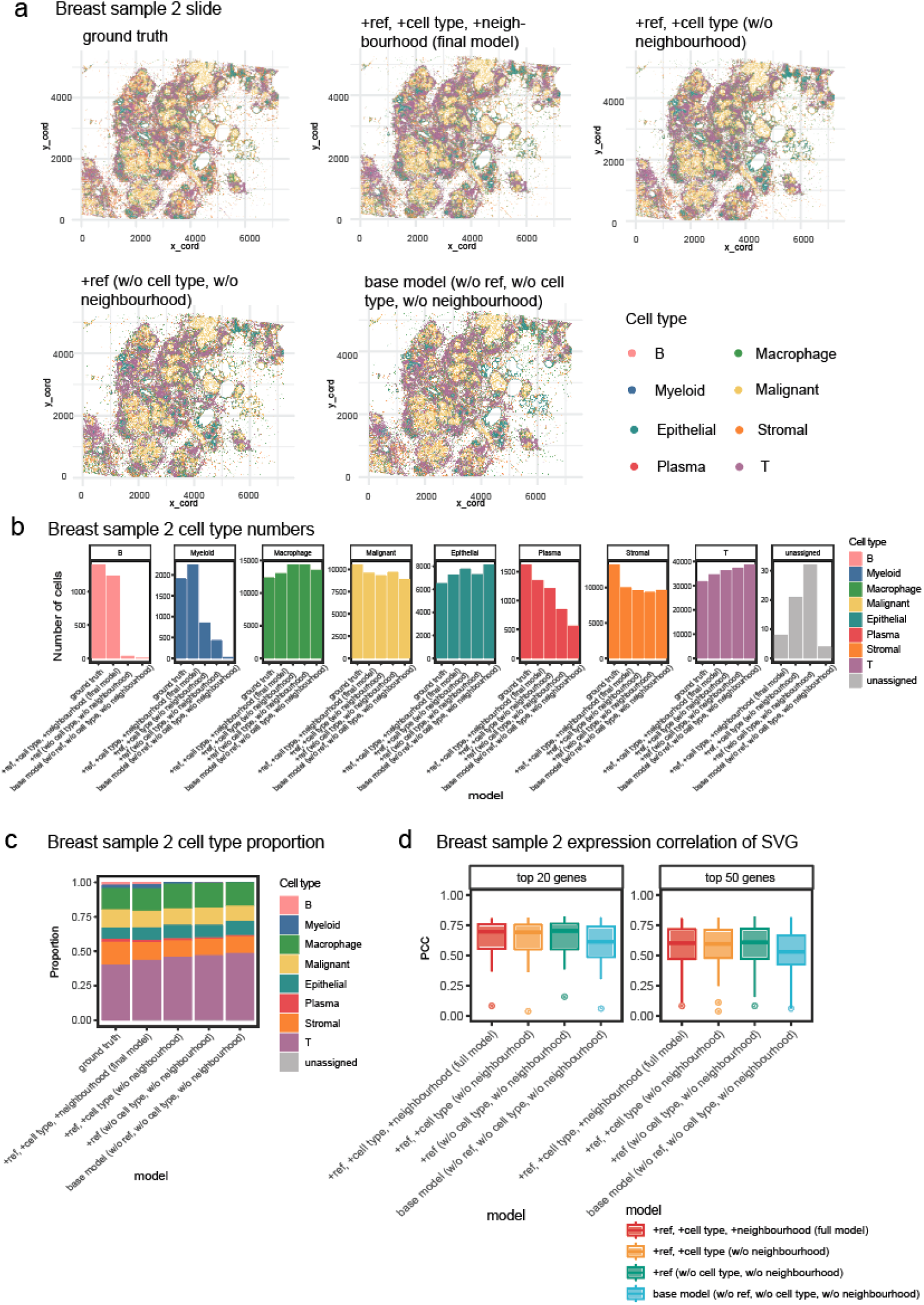
Ablation study. Performance of GHIST with and without key components (auxiliary cell type prediction, neighbourhood composition, and input single-cell reference data) on BreastCancer2. “w/o” stands for without the component, “+” represents with the components, “ref” stands for input single-cell reference data, “celltype” stands for auxiliary cell type prediction, “neighbourhood” stands for neighbourhood composition. **(a)** Cell types from scClassify using predicted single-cell expressions from the various settings mapped spatially across the slide. **(b)** Number of cells in each predicted cell type. B and myeloid cells were missing in predictions without auxiliary cell type or neighbourhood composition prediction. **(c)** Predicted cell type proportion. **(d)** PCC of predicted SVGs. PCC of the top 20 and top 50 SVGs were markedly lower in the base model without using single-cell reference data as input. Each boxplot ranges from the first to third quartile with the median as the horizontal line. The lower whisker extends 1.5 times the interquartile range below the first quartile, while the upper whisker extends 1.5 times the interquartile range above the third quartile.

**Supplementary Figure 2.**
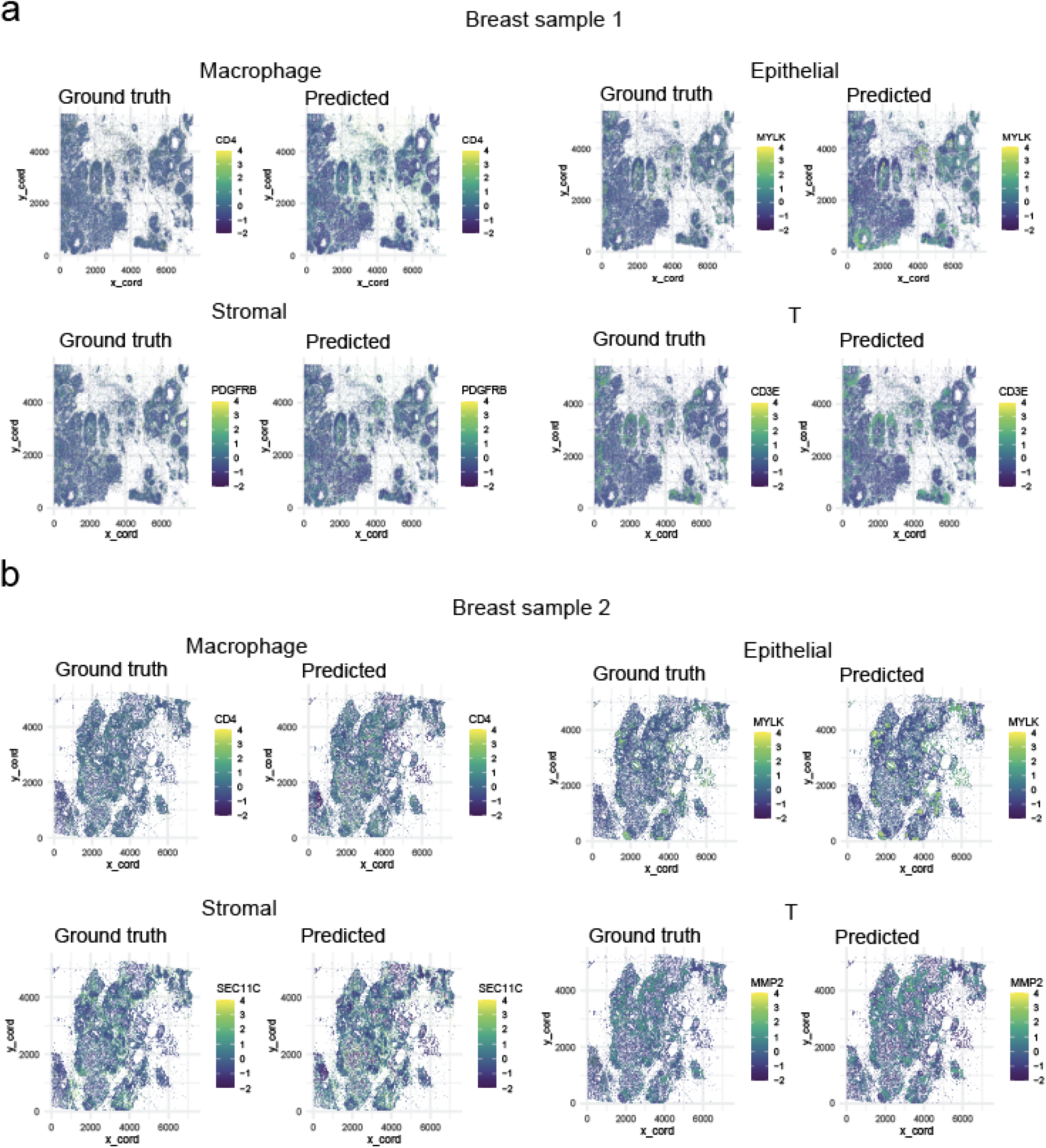
Additional markers of different cell types. Expression of predicted cell type markers (*CD4* for macrophages, *MYLK* for epithelial cells, *SEC11C* for stromal cells, and *MMP2* for T cells) mapped across the BreastCancer1 and BreastCancer2 slides.

**Supplementary Figure 3.**
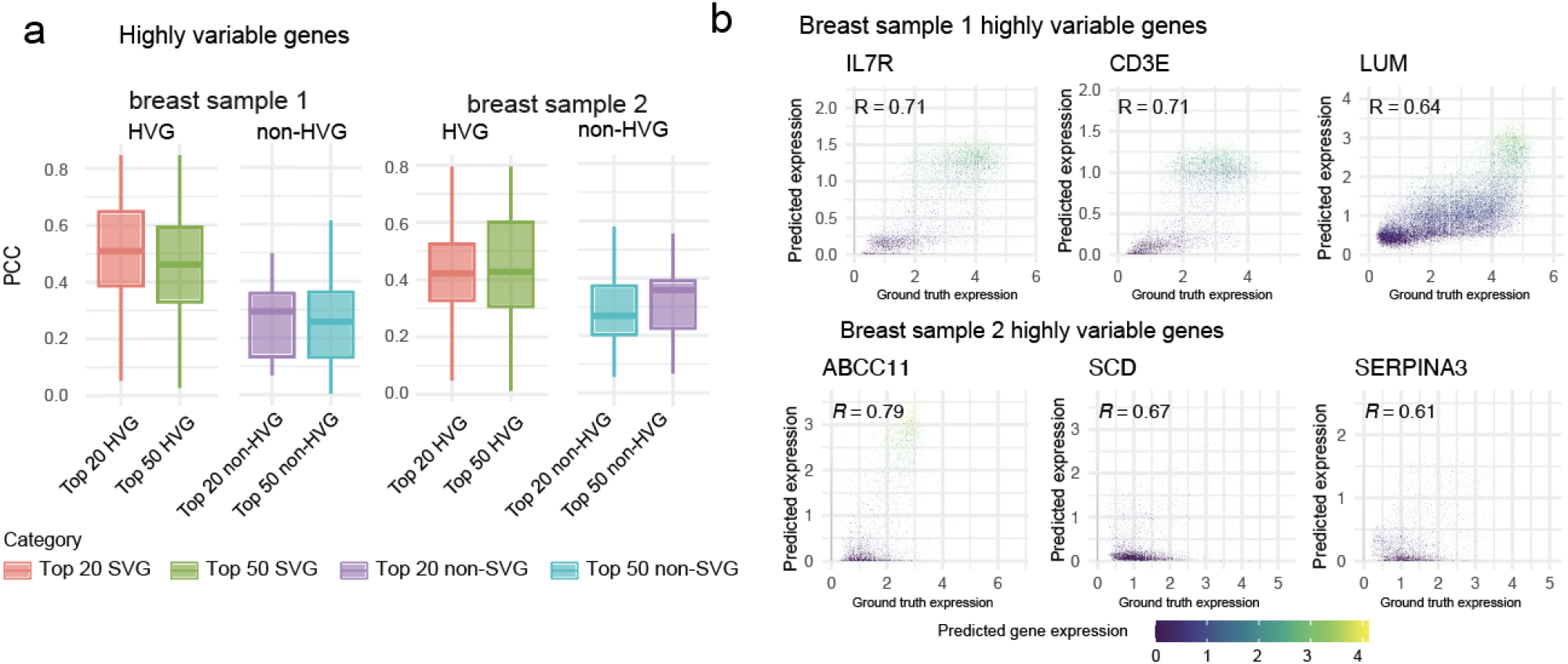
PCC of predicted HVGs in the two breast cancer Xenium samples. **(a)** Boxplots of the PCC for the top 20 and 50 genes between the ground truth and predicted expressions. Each boxplot ranges from the first to third quartile with the median as the horizontal line. The lower whisker extends 1.5 times the interquartile range below the first quartile, while the upper whisker extends 1.5 times the interquartile range above the third quartile. **(b)** Scatter plot between the ground truth and predicted expressions for selected HVGs (*IL7R, CD3E, LUM, ABCC11, SCD*, and *SERPINA3*) in BreastCancer1.

**Supplementary Figure 4.**
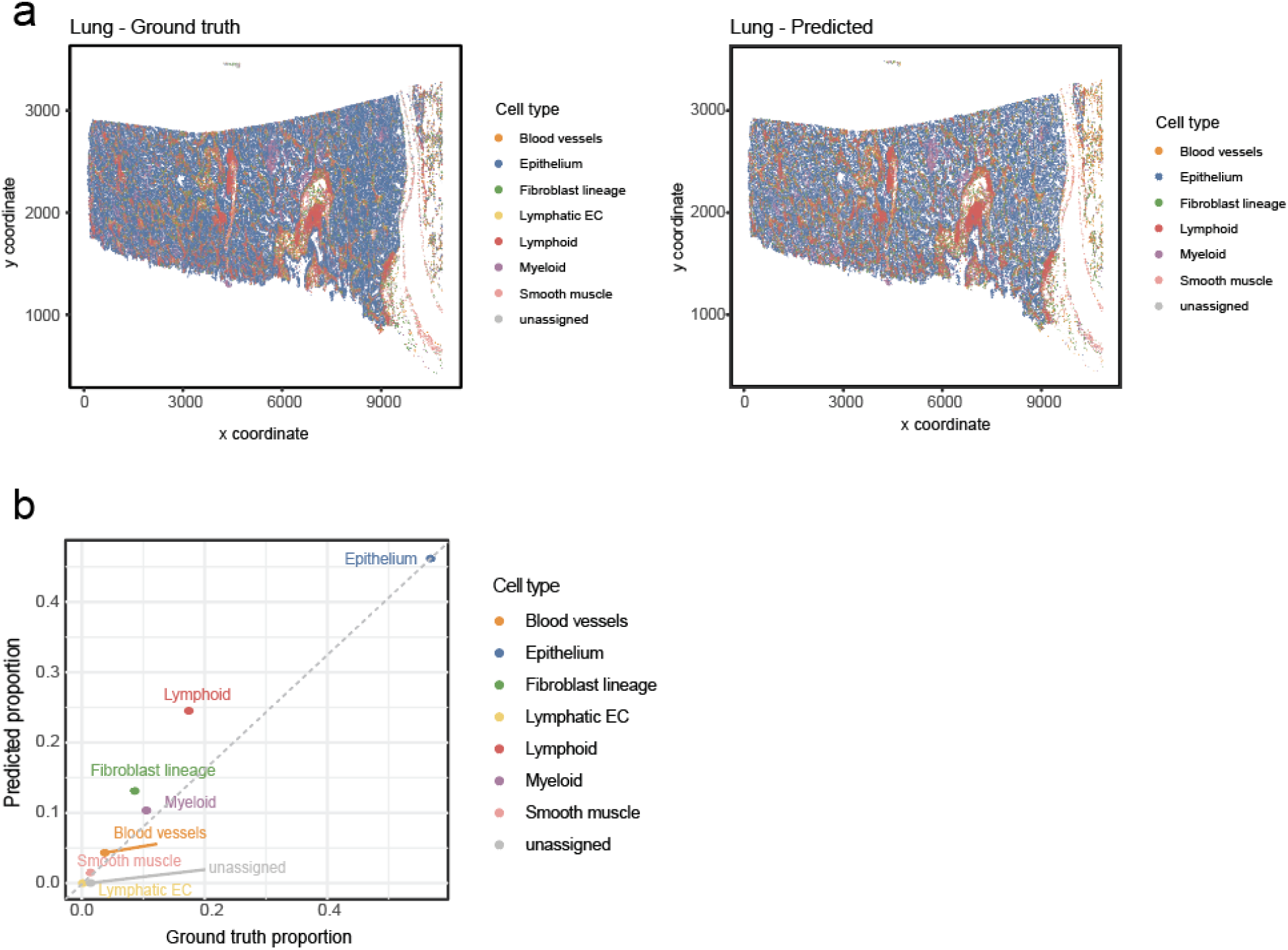
LungAdenocarcinoma results. **(a)** Cell types including blood vessels, epithelium, fibroblast lineage, lymphatic EC, lymphoid, myeloid, smooth muscle, and unassigned cells are mapped spatially across the tissue slide for the measured (ground truth) and predicted expressions. **(b)** Scatter plot between the measured (ground truth) and predicted cell type proportions.

**Supplementary Figure 5.**
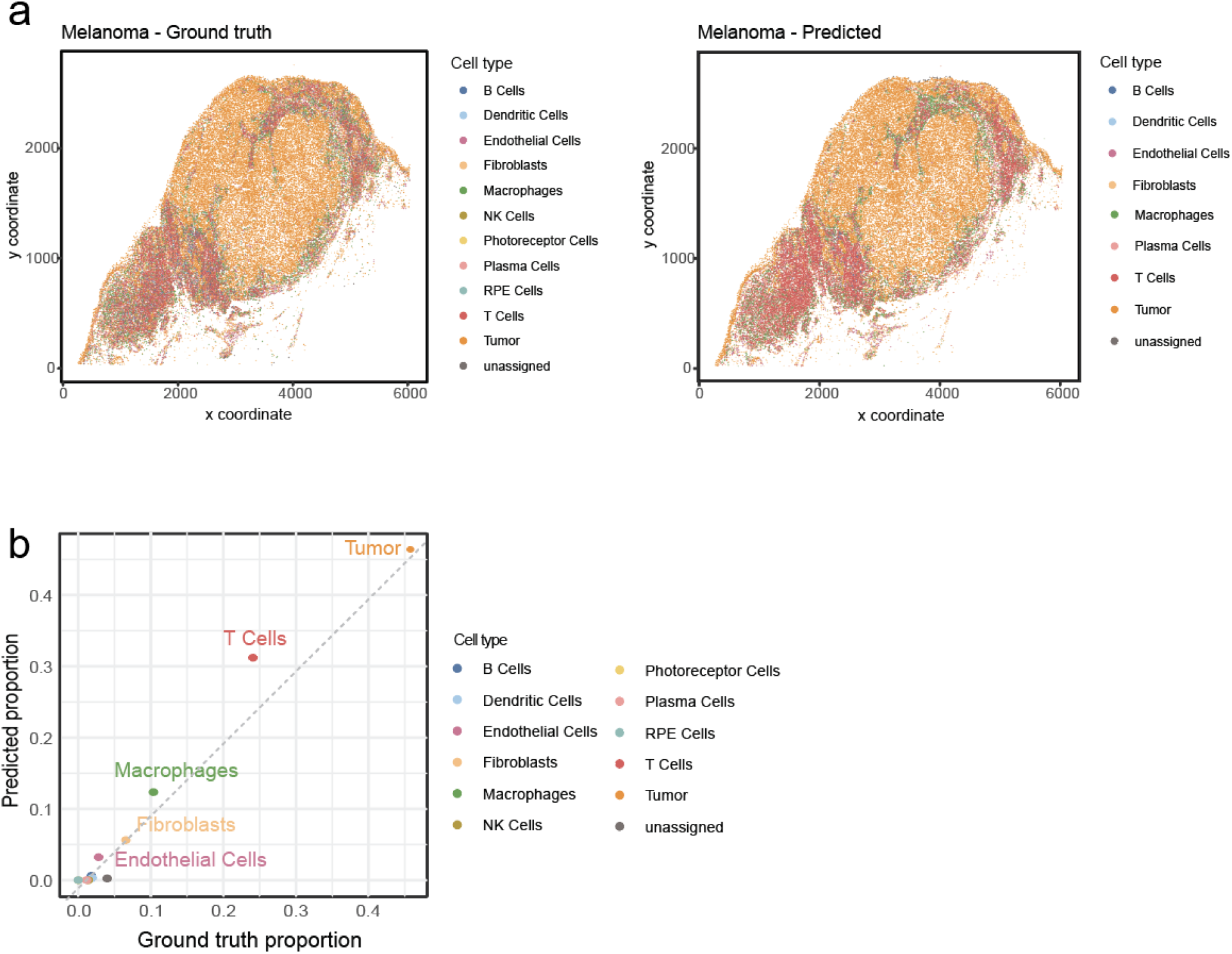
Melanoma results. **(a)** Cell types including B, dendritic, endothelial, fibroblasts, macrophages, NK, photoreceptor, plasma, RPE, T, tumour, and unassigned cells. These are mapped spatially across the tissue slide for the measured (ground truth) and predicted expressions. **(b)** Scatter plot between the measured (ground truth) and predicted cell type proportions.

**Supplementary Figure 6.**
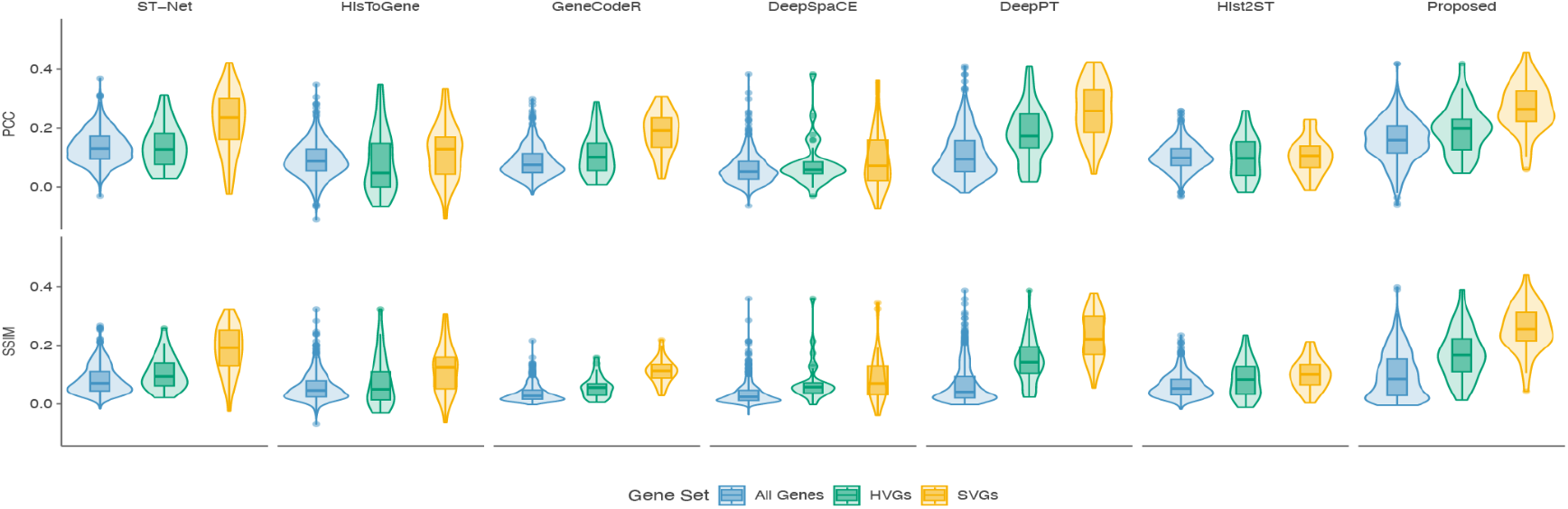
Spot-based gene prediction and survival analysis performance among state-of-the-art methods and GHIST using the HER2ST dataset. PCC and SSIM violin plots for each method for all genes as well as for selected HVGs and SVGs, demonstrating that selecting such types of genes is more biologically meaningful for comparison than all genes. Each plot ranges from the first to third quartile with the median as the horizontal line. The lower whisker extends 1.5 times the interquartile range below the first quartile, while the upper whisker extends 1.5 times the interquartile range above the third quartile.

**Supplementary Figure 7.**
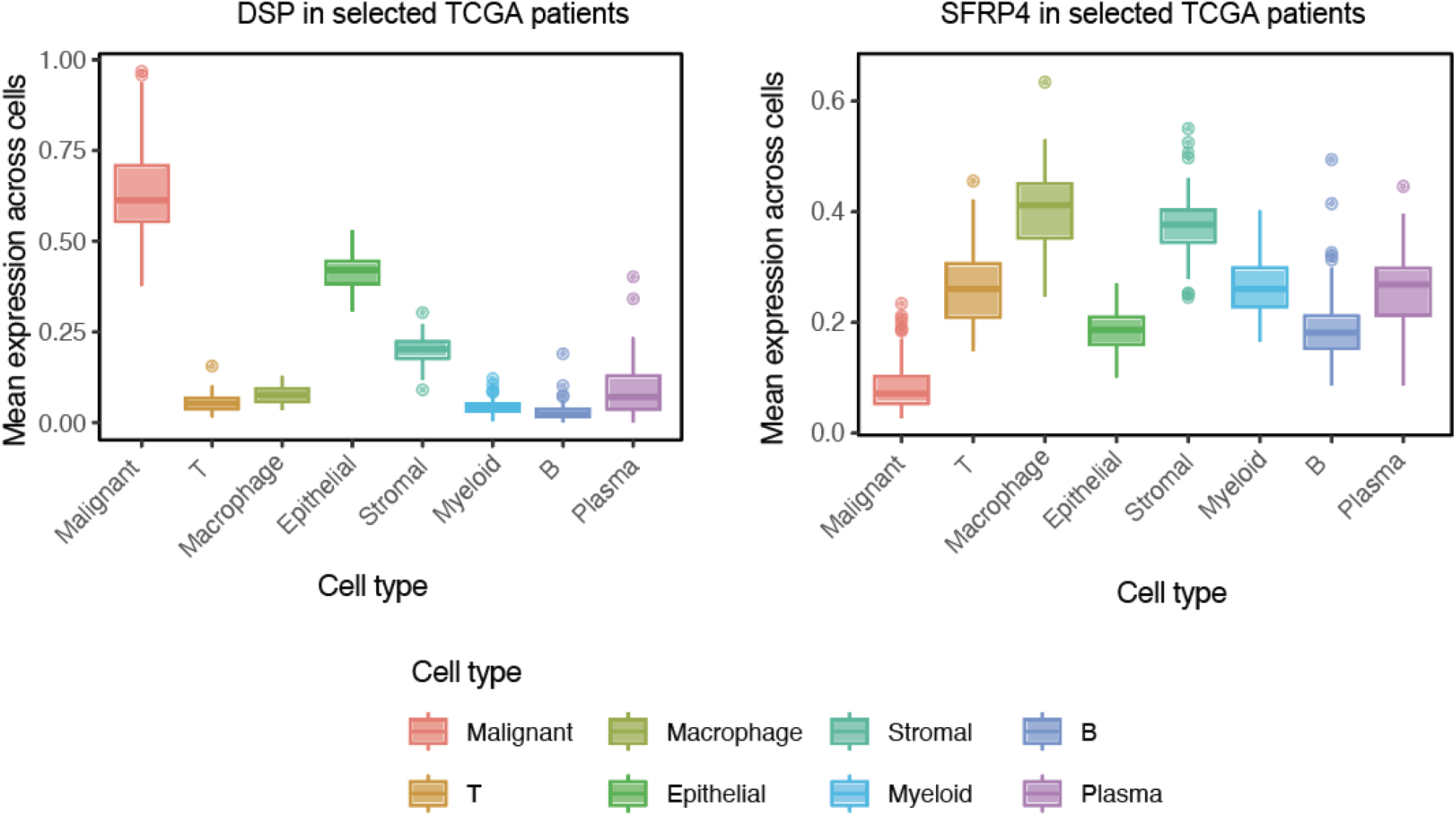
Predicted expression of selected genes in different cell types. Box plots showing the mean predicted expression of *DSP* and *SFRP4* across cells of different cell types in the TCGA patients. Each boxplot ranges from the first to third quartile with the median as the horizontal line. The lower whisker extends 1.5 times the interquartile range below the first quartile, while the upper whisker extends 1.5 times the interquartile range above the third quartile.

**Supplementary Figure 8.**
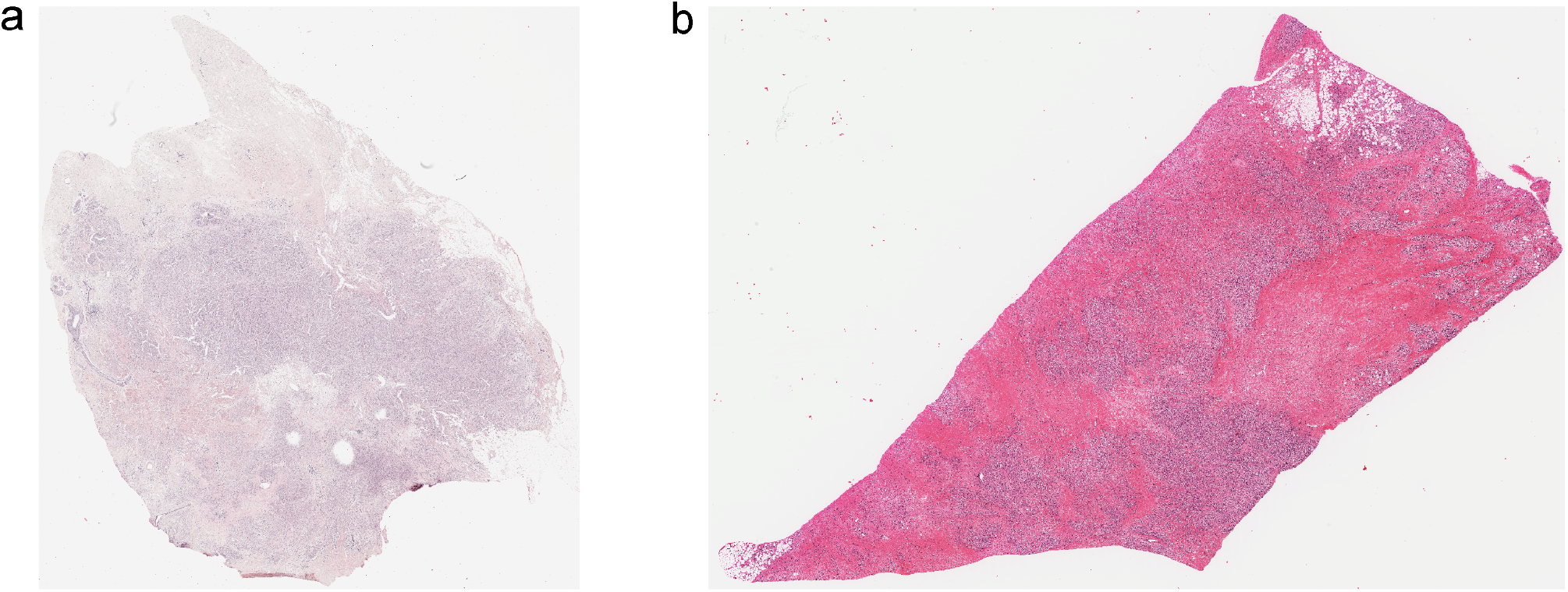
Stain quality issues in TCGA-BRCA. Examples of issues with stain quality observed in the H&E images from TCGA-BRCA patients, where **(a)** TCGA-AO-A12G appeared understained and faded, and **(b)** TCGA-PE-A5DD appeared very overstained.

**Supplementary Figure 9.**
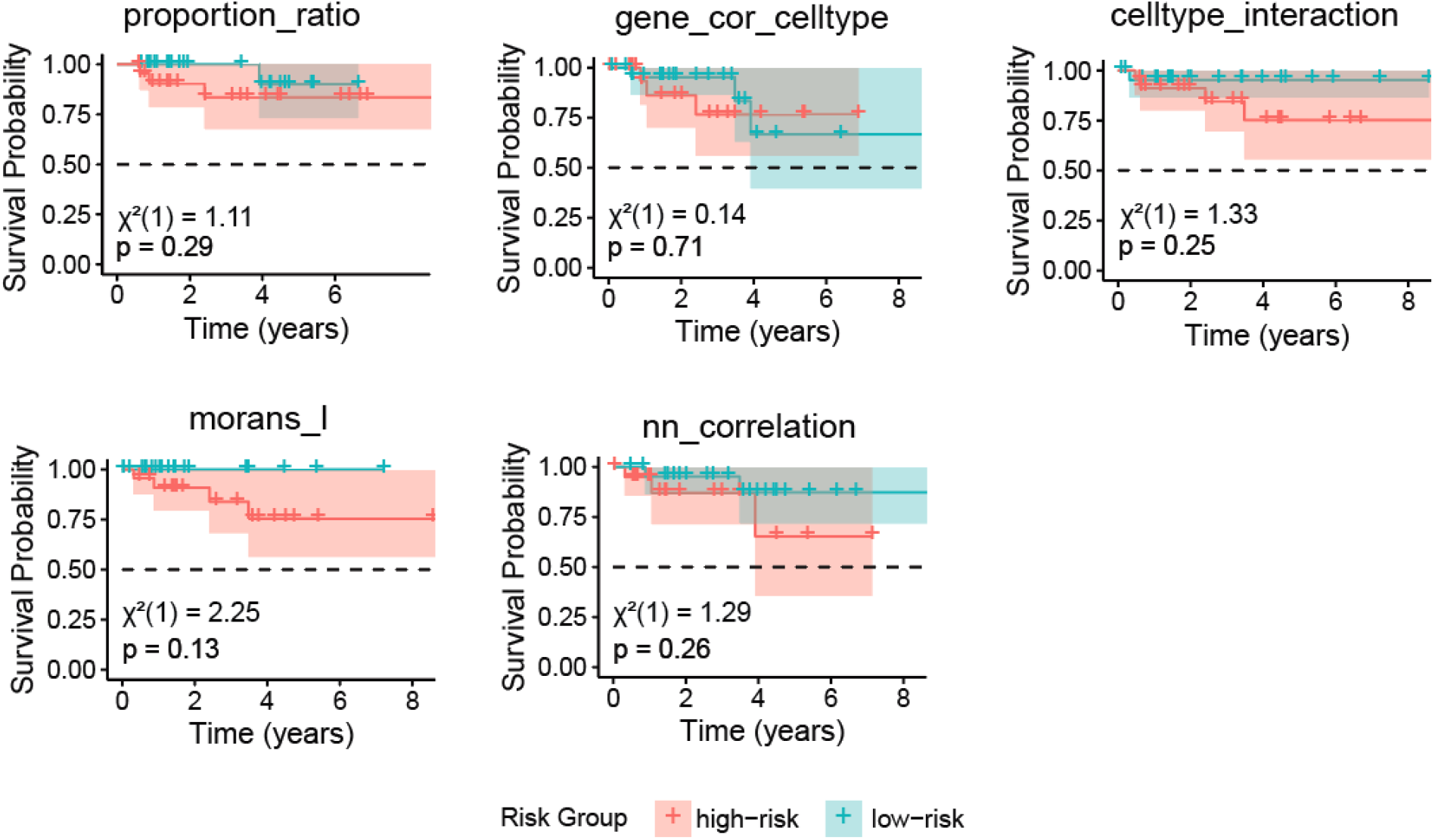
KM curves for the TCGA breast cancer HER2+ subset using various features types from scFeatures. Proportion_ratio represents the ratio between pairwise cell type proportions, gene_cor_celltype represents the correlation between pairwise gene expression, celltype_interaction represents the cell type composition of neighbouring cell types, morans_I represents the Moran’s I spatial autocorrelation metric, nn_correlation represents correlation of gene expression between neighbouring cells.

**Supplementary Figure 10.**
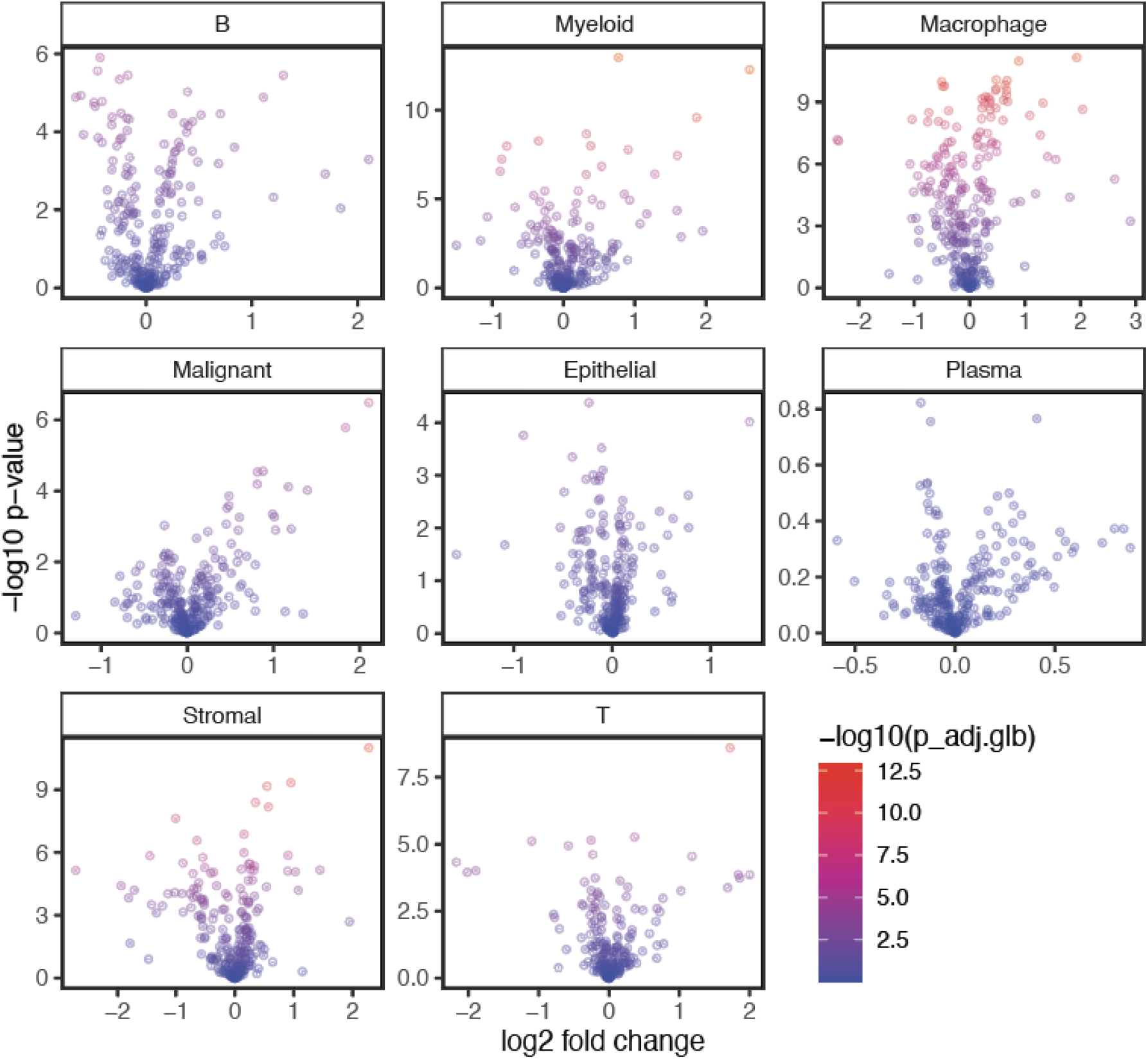
Volcano plots between Cluster 1 and Cluster 2 split by different cell types. Volcano plot showing significant differentially expressed genes between the cluster 1 and cluster 2 refined ER^+/^PR^+^ patients.

**Supplementary Figure 11.**
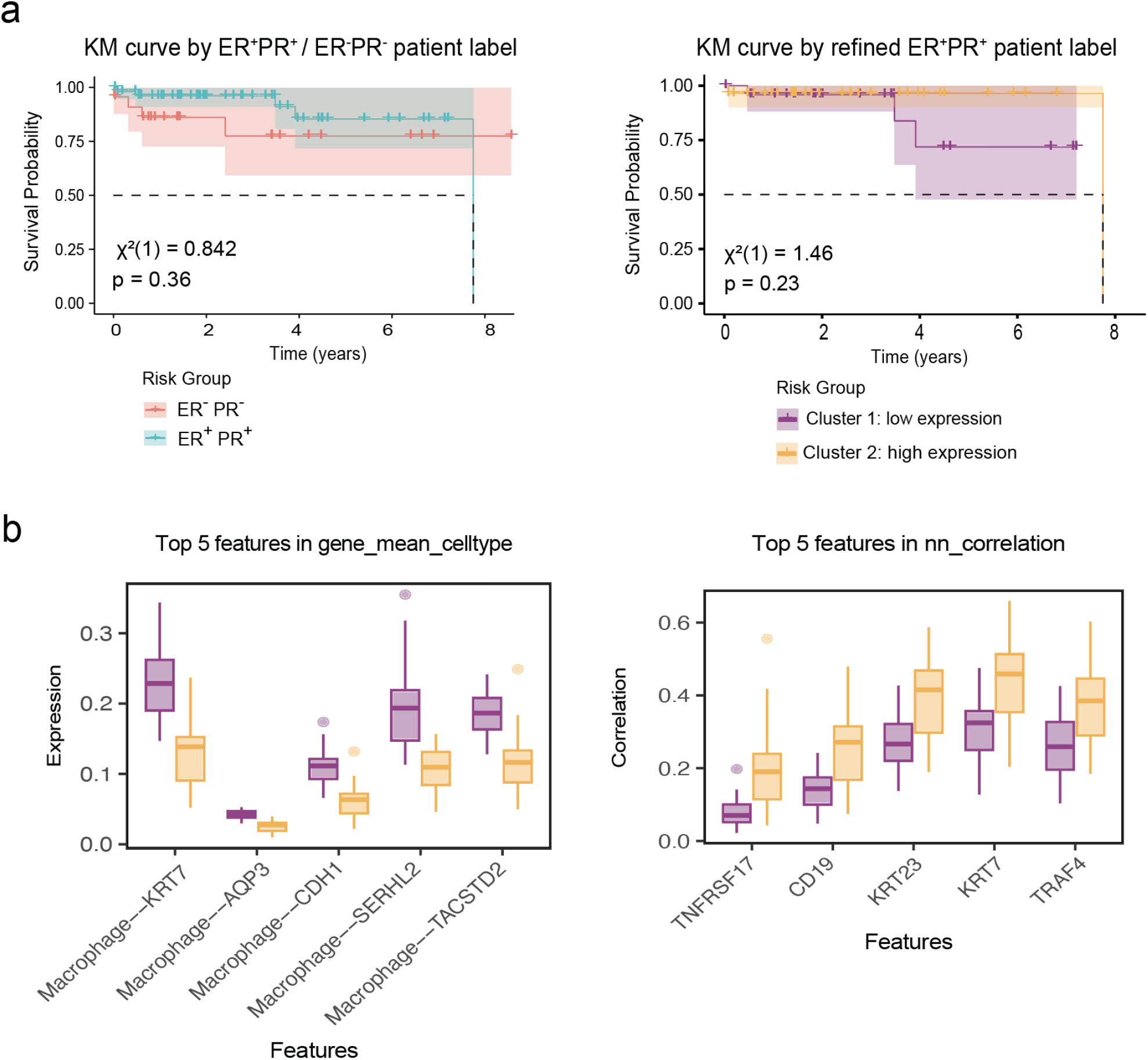
KM curves of original patient label and refined patient label. **(a)** The KM curve based on the original ER+/PR+ and ER-/PR- patient group as defined in TCGA [left], and the KM curve based on refined patient clusters within the ER+PR+ patients [right]. **(b)**, the top 5 features with greatest feature importance in the cluster 1 and cluster 2 classification model based on the cell-type-specific gene mean expression feature type [left], and the top 5 features in the model based on the nearest neighbour correlation feature type [right]. Each boxplot ranges from the first to third quartile with the median as the horizontal line. The lower whisker extends 1.5 times the interquartile range below the first quartile, while the upper whisker extends 1.5 times the interquartile range above the third quartile.

## Notes

### Competing Interest Statement

The authors have declared no competing interest.

